# Landscape drives zoonotic malaria prevalence in non-human primates

**DOI:** 10.1101/2022.06.17.496284

**Authors:** Emilia Johnson, Reuben Sunil Kumar Sharma, Pablo Ruiz Cuenca, Isabel Byrne, Milena Salgado-Lynn, Zarith Suraya Shahar, Lee Col Lin, Norhadila Zulkifli, Nor Dilaila Mohd Saidi, Chris Drakeley, Jason Matthiopoulos, Luca Nelli, Kimberly Fornace

**Affiliations:** School of Biodiversity, One Health and Veterinary Medicine, University of Glasgow, G12 8QQ, Scotland; Department of Disease Control, London School of Hygiene & Tropical Medicine, London WC1E 7HT, UK; Centre on Climate Change and Planetary Health, London School of Hygiene & Tropical Medicine, London WC1E 7HT, UK; Faculty of Veterinary Medicine, Universiti Putra Malaysia, 43400, UPM, Serdang, Selangor, Malaysia; Lancaster University, Bailrigg, Lancaster LA1 4YW; Liverpool School of Tropical Medicine, Pembroke Place Liverpool, L3 5QA, UK; School of Biosciences, Cardiff University, Cardiff CF10 3AX, UK; Wildlife Health, Genetic and Forensic Laboratory, Sabah Wildlife Department, Wisma Muis, 88100, Kota Kinabalu, Malaysia; Danau Girang Field Centre, Sabah Wildlife Department, Wisma MUIS 88100, Kota Kinabalu Sabah, Malaysia; Department of Infection Biology, London School of Hygiene & Tropical Medicine, London WC1E 7HT, UK; Saw Swee Hock School of Public Health, National University of Singapore, Singapore 117549

**Keywords:** Disease ecology, habitat fragmentation, landscape change, malaria, *Plasmodium knowlesi*, zoonoses

## Abstract

Zoonotic disease dynamics in wildlife hosts are rarely quantified at macroecological scales due to the lack of systematic surveys. Non-human primates (NHPs) host *Plasmodium knowlesi,* a zoonotic malaria of public health concern and the main barrier to malaria elimination in Southeast Asia. Understanding of regional *P. knowlesi* infection dynamics in wildlife is limited. Here, we systematically assemble reports of NHP *P. knowlesi* and investigate geographic determinants of prevalence in reservoir species. Meta-analysis of 6322 NHPs from 148 sites reveals that prevalence is heterogeneous across Southeast Asia, with low overall prevalence and high estimates for Malaysian Borneo. We find that regions exhibiting higher prevalence in NHPs overlap with human infection hotspots. In wildlife and humans, parasite transmission is linked to land conversion and fragmentation. By assembling remote sensing data and fitting statistical models to prevalence at multiple spatial scales, we identify novel relationships between *P. knowlesi* in NHPs and forest fragmentation. This suggests that higher prevalence may be contingent on habitat complexity, which would begin to explain observed geographic variation in parasite burden. These findings address critical gaps in understanding regional *P. knowlesi* epidemiology and indicate that prevalence in simian reservoirs may be a key spatial driver of human spillover risk.

## INTRODUCTION

Zoonotic infectious diseases arise from the spillover of pathogens into human populations, typically from a reservoir in wildlife hosts. Anthropogenic land use and land cover change have now been widely linked to infectious disease outbreaks (Brock et al., 2019; Davidson et al., 2019a; Loh et al., 2016). Such practices, including deforestation, logging, clearing for cash-crop plantations or conversion of intact forest into arable land, are accelerating across tropical forests of Southeast Asia (Fornace et al., 2021; Imai et al., 2018)(Fornace et al., 2021; Imai et al., 2018). Mechanisms that underly the association between habitat disturbance and spillover risk from wildlife hosts are complex and occur over multiple spatial scales (Brock et al., 2019). In Brazil, re-emergence of Yellow Fever Virus in both NHPs and humans has been linked to areas with highly fragmented forest (Ilacqua et al., 2021). In part, an increase in ‘edge’ habitat in fragmented or mosaic landscapes can facilitate spatial overlap and altered contact patterns between wildlife, vectors and humans (Lehman et al., 2006). Such ecological interfaces are also thought to contribute to parasite spillover in other vector-borne diseases including Zika (J. Li et al., 2021), Babesiosis and Lyme disease (Simon et al., 2014), *Trypanosoma cruzi* (Vaz et al., 2007) and zoonotic malaria (Brock et al., 2019; Grigg et al., 2017). At the same time, habitat fragmentation can have detrimental impact on wildlife population viability, with reduced host species occupancy and reduced disease burden in highly disturbed habitats (Hanski and Ovaskainen, 2000). Disentangling this interplay is essential to inform ecological strategies for surveillance and mitigation of diseases in regions undergoing landscape change (Fornace et al., 2021).

Zoonotic *P. knowlesi* is a public health threat of increasing importance across Southeast Asia, following the identification of a prominent infection foci in Borneo in 2004 (Singh et al., 2004). *P. knowlesi* is a zoonosis, with a sylvatic cycle circulating in non-human primates (NHPs). Human cases currently occur only from spillover events (Cuenca et al., 2022; Fornace, 2022; Fornace et al., 2023; Lee et al., 2011). Human transmission requires bites from infective mosquitos, primarily anopheline mosquitos of the Leucosphyrus Complex (*Anopheles balabacensis, An. latens, An. introlactus)* and Dirus Complex (*An. dirus, An. cracens)* (Moyes et al., 2016; Vythilingam et al., 2006; Wong et al., 2015). Natural hosts for *P. knowlesi* are typically Long-tailed macaques (*Macaca fascicularis*) and Southern Pig-tailed macaques (*M. nemestrina)* (Moyes et al., 2016), both occurring widely across Southeast Asia. Currently, distribution of *P. knowlesi* cases is thought to be restricted to the predicted ranges of known vector and host species (Davidson et al., 2019b), though recent studies have also identified other NHPs found to be harbouring *P. knowlesi.* This includes Stump-tailed macaques *(M. arctoides),* which are now considered to be another natural reservoir (Fungfuang et al., 2020).

Progress towards malaria elimination in Malaysia has been stymied by a recent rise in human incidence of *P. knowlesi* malaria. Even after accounting for increases in surveillance and diagnostic improvements it is now recognised as the most common cause of clinical malaria in Malaysia (Cooper et al., 2020). Indeed, Malaysia was the first country not to qualify for malaria elimination due to ongoing presence of zoonotic malaria and the World Health Organisation (WHO) updated the guidelines to reflect zoonotic malaria as a public health threat (“Global Malaria Programme,” n.d.). Emergence of *Plasmodium knowlesi* infections has been linked to changes in land cover and land use (Fornace et al., 2021). While sporadic cases have been reported across Southeast Asia, including in Indonesia (Setiadi et al., 2016), the Philippines (Fornace et al., 2018), Vietnam (Maeno et al., 2015), Brunei (Koh et al., 2019) and Myanmar (Ghinai et al., 2017), the majority of *P. knowlesi* cases are found in East Malaysia (Borneo) with hotspots in the states of Sabah and Sarawak (Jeyaprakasam et al., 2020), areas that have seen extensive deforestation and landscape modification. In Sabah, human prevalence of *P. knowlesi* infection has recently been shown to be specifically associated with recent loss of intact forest, agricultural activities, and fragmentation across multiple localised spatial scales (Brock et al., 2019; Fornace et al., 2019b, 2016).

Prevalence of the pathogen in reservoir hosts is one of three crucial factors determining the force of infection in zoonotic spillover events (Murray and Daszak, 2013). Despite this, very little is known of the impact of rapid landscape change on the distribution of *P. knowlesi* in NHPs. Literature on the impacts of fragmentation on primates tends to focus on primate density and abundance (Link et al., 2010; ZUNINO et al., 2007). What is known is that effects of land cover changes on primate-pathogen dynamics are highly variable and context-specific. Although the vector species responsible for sylvatic transmission remain unknown, the *Anopheles leucospryphus* group, the only vector group implicated in *P. knowlesi* transmission, is widely associated with secondary, disturbed forest (Brant, 2011; Hawkes et al., 2019; Wong et al., 2015). Macaques have been known to preferentially rely on fringe habitat, a behaviour that may be exaggerated in response to habitat fragmentation and facilitate exposure to vectors (Lehman et al., 2006; Stark et al., 2019). Changes to land composition can also create the biosocial conditions for higher rates of parasitism in primates. Under conditions of limited resources and reduction in viable habitat, conspecific primate density may increase as troops compete for available space. In turn, this can favour transmission via intra-species contact or allow the exchange of pathogens between troops dwelling in interior forest versus edge habitat (Faust et al., 2018; Stark et al., 2019). Habitat use may also become more intensive, preventing parasite avoidance behaviours (Nunn and Dokey, 2006). Land cover change is also known to favour more adaptable, synanthropic species such as *M. fascicularis* (McFarlane et al., 2012). Considering the spillover risk posed by wildlife reservoirs of *P. knowlesi,* clarifying any relationships between environmental factors and parasitaemia in key host species may contribute to a more comprehensive understanding of *P. knowlesi* transmission patterns.

Earth Observation (EO) data provides novel opportunities to investigate epidemiological patterns of diseases which are linked to environmental drivers (Kalluri et al., 2007). In relation to *P. knowlesi,* utility of fine-scale remote-sensing data has been demonstrated: examples include satellite-derived data used to examine household–level exposure risk in relation to proximate land configuration (Fornace et al., 2019b), UAV-imagery used to link real–time deforestation to macaque host behavioural change (Stark et al., 2019), and remote-sensing data used to interrogate risk factors for vector breeding sites (Byrne et al., 2021). Though macroecological studies that utilise geospatial data are often confounded by issues of matching temporal and spatial scales, as well as by the quality and accuracy of available georeferencing, measures can be taken to account for this when examining the role of environmental factors in modulating disease outcomes. Furthermore, ecological processes occur and interact over a range of distances, or ‘spatial scales’ (Brock et al., 2019; Fornace et al., 2016; Loh et al., 2016). This applies to determinants of vector-borne disease ecology, from larval breeding microclimate to wildlife host foraging behaviour. As multiple influential variables are rarely captured by a single scale (Cohen et al., 2016), data-driven methods can be applied to examine risk factors over multiple scales and identify covariates at their most influential extent (Byrne et al., 2021).

We hypothesise that prevalence of *P. knowlesi* in primate host species is spatially heterogeneous and that higher prevalence is partially driven by forest loss and fragmentation, contributing to the strong associations described between land use, land cover and human *P. knowlesi* risk. This study is the first to systematically assess *P. knowlesi* prevalence in NHPs at a regional scale, and across a wide range of habitats. In conceptual frameworks and transmission models, it is often assumed that *P. knowlesi* infections in NHPs are chronic (low level, persistent infection) and ubiquitous (uniformly distributed across populations) (Brock et al., 2016; Jeyaprakasam et al., 2020). No studies have systematically assessed the extent and quality of all available data on *P. knowlesi* in NHPs. Independent studies investigating *P. knowlesi* in primates are typically constrained by small sample sizes and confined geographic areas, limiting inference that can be made about relationships between infection dynamics and landscape characteristics. Systematic tools developed for epidemiological studies of disease prevalence in human populations are rarely applied to the study of wildlife disease prevalence; however, such tools can be used to capture the scale and contrast required in macroecological studies to quantify disease burdens regionally. Furthermore, while recent research has shown the impact of deforestation on the distribution of macaques in the context of *P. knowlesi* (Moyes et al., 2016; Stark et al., 2019), associations between landscape and variation in the prevalence of simian *Plasmodium* spp. in primates have not been explored. We aimed to 1) assemble a georeferenced dataset of *P. knowlesi* in NHPs; 2) evaluate variation in NHP *P. knowlesi* prevalence by geographic region; and 3) assess environmental and spatial risk factors for *P. knowlesi* prevalence in NHPs across Southeast Asia.

## RESULTS

A systematic literature review was conducted in Medline, Embase and Web of Science to identify articles reporting prevalence of naturally acquired *Plasmodium knowlesi* in NHPs. 23 research articles were identified (Akter et al., 2015; Amir et al., 2020; Chang et al., 2011; Fungfuang et al., 2020; Gamalo et al., 2019; Ho et al., 2010; Jeslyn et al., 2011; Lee et al., 2011; M. I. Li et al., 2021; Muehlenbein et al., 2015; Putaporntip et al., 2010; Saleh Huddin et al., 2019; Seethamchai et al., 2008; Unpublished, 2015, 2013; Vythilingam et al., 2008; Zarith et al, 2021; Zhang et al., 2016), containing 148 unique primate survey records to form the dataset for analyses (see SI for details of JBI Critical Assessment, Table S5) (Munn et al., 2015). Year of sampling ranges from 2004–2019. No primatological studies were identified from Vietnam, Brunei or Timor-Leste. Full characteristics of the articles and individual study methodologies are reported in Supplementary Information (Table S2). Spatial resolution of the survey sites varied from GPS point coordinates to country-level administrative boundaries (Supplementary Table S7). Geographic distribution of sampling is illustrated in Figure 1.

**Figure 1.**
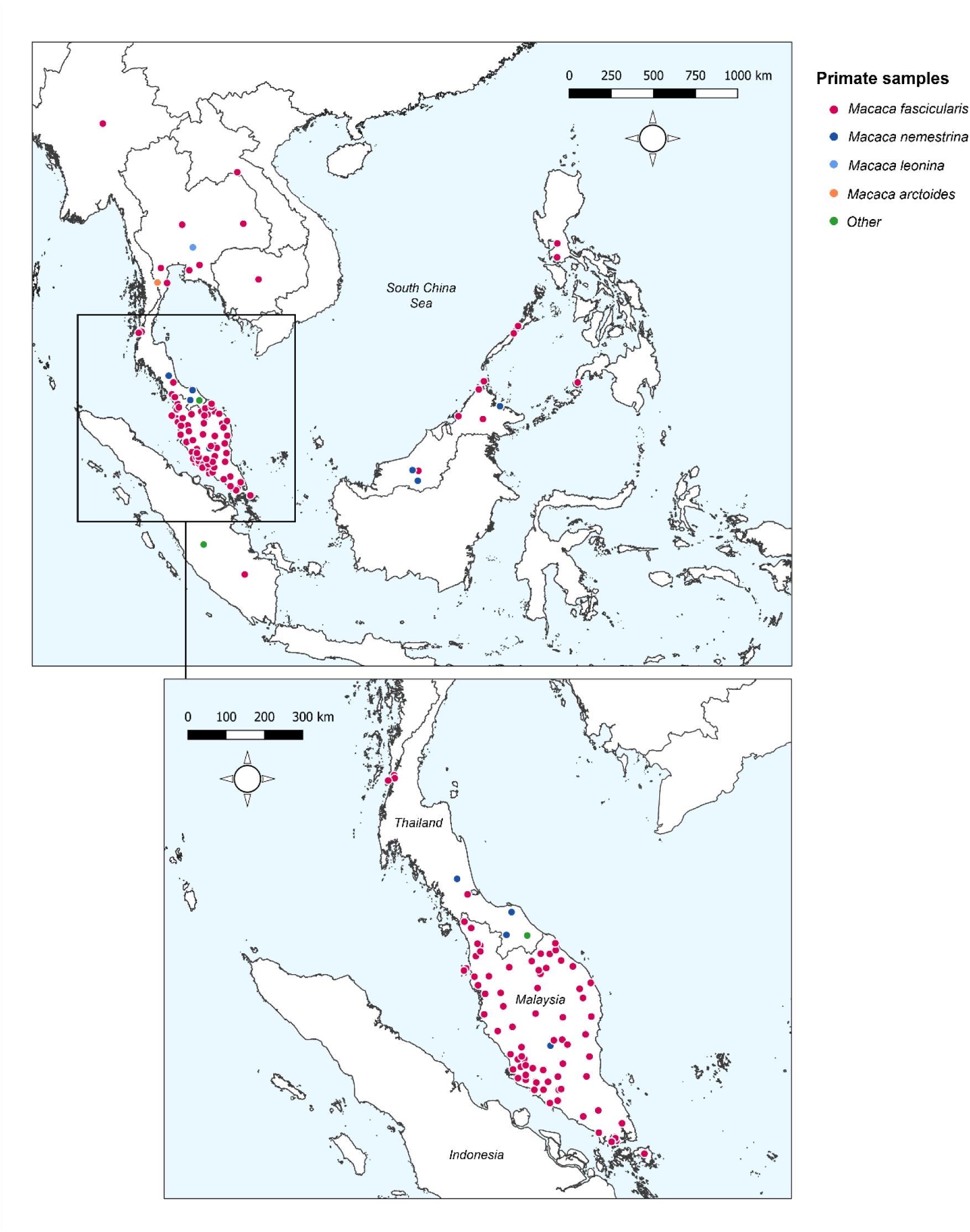
Sampling sites and primate species sampled across Southeast Asia. ‘Other’ includes Trachypithecus obscurus *and undefined species from the genus* Presbytis. *Total surveys = 148*.

Overall, records report on a total of 6322 primates, with the largest proportion sampled from Peninsular Malaysia (48.5%, n=3069/6322). Primate surveys were primarily conducted on Long-tailed macaques (*Macaca fascicularis*) (90.5%, n=5720/6322) followed by Pig-tailed macaques (*M. nemestrina)* (n=532/6322) (Amir et al., 2020; Lee et al., 2011; Muehlenbein et al., 2015; Putaporntip et al., 2010) (Table S3). Reported prevalence of *Plasmodium knowlesi* in NHPs ranged from 0%–100%. Only 87 of the surveys (58.8%, n=87/148) reported a positive diagnosis, with the remaining 61 sites finding no molecular evidence of *P. knowlesi* infection (41.2%) in any primates tested. A full breakdown of *P. knowlesi* infection rates according to reported primate characteristics can be found in SI, Table S4.

### Meta-analysis of *P. knowlesi* prevalence

To quantify regional heterogeneity in simian cases of *P. knowlesi,* a one-stage meta-analysis of prevalence (number positive out of the number sampled) was conducted on primate malaria survey data. Overall pooled estimate for *P. knowlesi* prevalence was 11.99% (CI95% 9.35–15.26). Overall heterogeneity was assessed using the I^2^ statistic. Substantial between-study heterogeneity (I^2^ ≥75%) was found across all prevalence records (I^2^=80.5%; CI95% 77.3–83.1). In the sub-group analysis by region, pooled prevalence estimates are consistently low for Thailand (2.0%, CI95% 1.1–3.5%), moderate in Peninsular Malaysia (14.3%, CI95% 11.1–18.2) and elevated in Singapore (23.3%, CI95% 11.0–42.8) and Malaysian Borneo (41.1%, CI95% 20.8–64.9) (Figure 2). Sub-group heterogeneity was assessed using prediction intervals, derived from τ ^2^ statistic used to describe between-study variability. Prediction intervals indicate high heterogeneity of estimates within regions, consistent with expectations of high variability of prevalence across individual study sites. Detailed forest plots for individual prevalence estimates can be found in Supplementary Figures S6.

**Figure 2.**
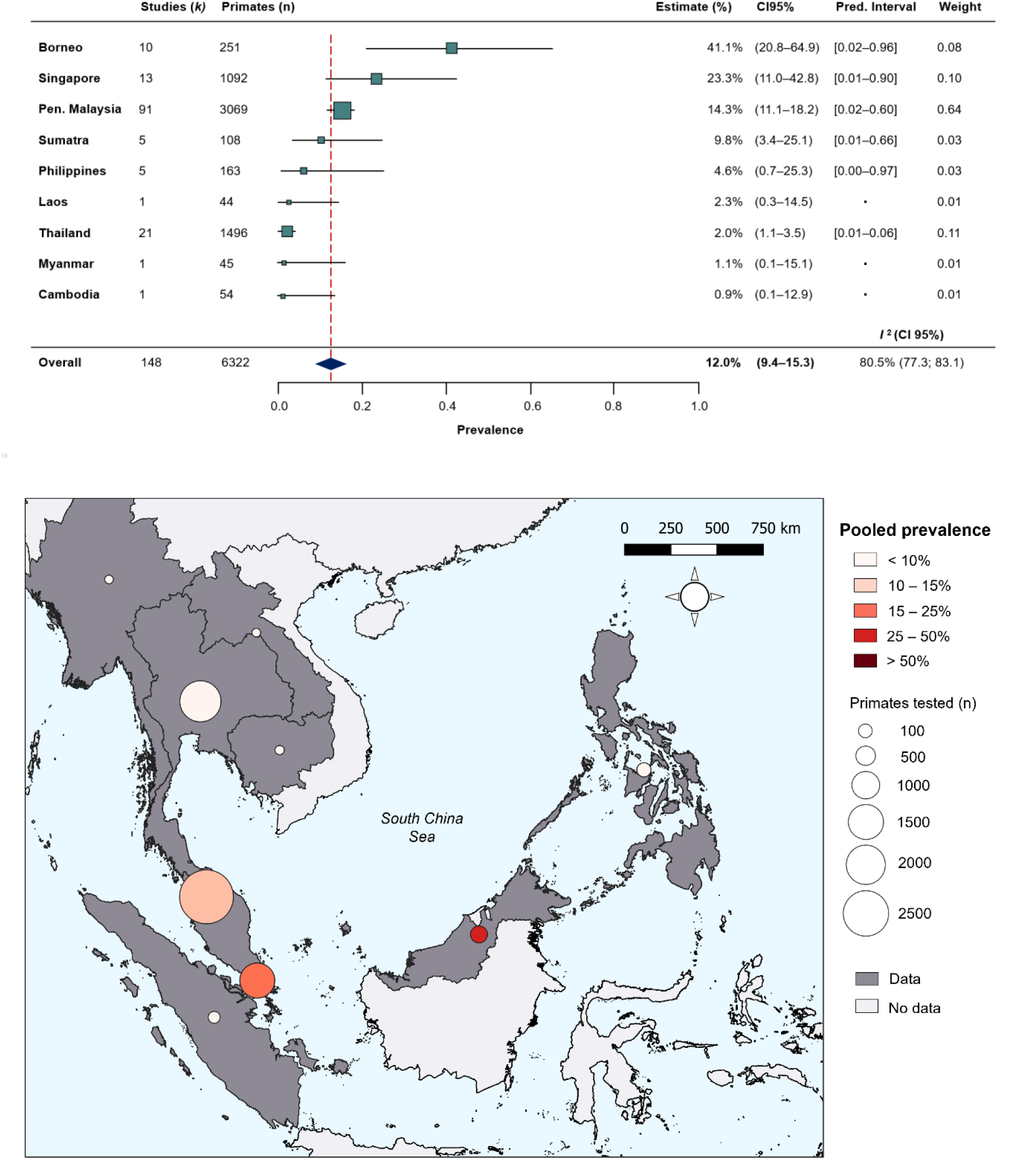
**(A)** Forest plot of pooled estimates for P. knowlesi prevalence (%) in all non-human primates tested (n=6322) across Southeast Asia, disaggregated by species and sampling site (k=148). Random-effects meta-analysis, sub-grouped by region. **(B)** Map of regional prevalence estimates for P. knowlesi prevalence in NHP in Southeast Asia from meta-analysis. Point colour denotes pooled estimate (%). Size denotes total primates tested per region (n). Shading indicates data availability.

### Risk factor analysis

Covariate data and *P. knowlesi* prevalence data were used to fit additional models to explore the relationships between localised landscape configuration and NHP malaria prevalence. Environmental covariates were extracted from satellite-derived remote sensing datasets (Table 1) at either true sampling sites (GPS coordinates) or 10 random pseudo-sampling sites to account for geographic uncertainty in prevalence data. Host species was grouped as ‘*Macaca fascicularis’* or ‘Other’ due to sample counts of <10 for certain primate species. Only 57.4% (n=85/148 records) of data included year of sampling, deemed to be insufficient to assess temporal patterns in prevalence. Tree canopy cover ranged from negligible to near total cover (100%) within buffer radii (Supplementary Table S12). Details of covariate data processing is illustrated in Supplementary Information (Figure S7–8).

**Table 1.** Spatial and temporal resolution and sources for environmental covariates. Summary metrics extracted within 5, 10 and 20km circular buffers.

| Covariate | Spatial res. | Temporal res. | Source |
| --- | --- | --- | --- |
| Human density (p/km<sup>2</sup>) | 1km | 2012 | WorldPop (WorldPop, 2018) |
| Elevation (m) | 1km | 2003 | SRTM 90m Digital Elevation v4.1 (Jarvis, A., H.I. Reuter, A. Nelson, 2008) |
| Tree cover (1/0) | 30m | Annual | Hansen’s Global Forest Watch (Hansen et al., 2013) |
| *Derivatives* | Proportion canopy cover (%) |  |  |
|  | Perimeter: area ratio (PARA>0) |  |  |

Following a two-stage approach for selection of explanatory variables, tree cover and fragmentation (measured by perimeter: area ratio, PARA) were retained at 5km as linear terms, human population density was retained at both 5km and 20km and primate species was retained as a categorical variable. Spearman’s rank tests for residual correlation between final variables at selected scales indicates a strong negative correlation between tree cover and fragmentation index (PARA) (ρ= –0.75) (SI, Figure S14).

Adjusting for all other covariates in the model, we identified strong evidence of an effect between increasing tree canopy cover and higher prevalence of *P. knowlesi* in NHPs within a 5km radius (aOR=1.38, CI95% 1.19–1.60; *p*<0.0001). Evidence was also found for an association between likelihood of *P. knowlesi* and higher degrees of habitat fragmentation (PARA) within 5km (aOR=1.17, CI95% 1.02–1.34, *p*<0.0281). Evidence suggests that human population density within a 5km radius is associated with risk of *P. knowlesi* in NHP (aOR=1.36, CI95% 1.16–1.58, *p=*0.0001) whilst human density within 20km has an inverse effect on likelihood of *P. knowlesi* (aOR=0.56, CI95% 0.46–0.67, *p*<0.0001). *M. fascicularis* is also associated with higher prevalence relative to all other non-human primate species (aOR=2.50, CI95% 1.31–4.85; *p*=0.0051). Additional complexity did not improve optimal model fit and effect modification was not pursued. In sensitivity analyses removing data points with excessive spatial uncertainty or restricting data points only to areas with high probability of macaque occurrence, evidence was consistently found that tree canopy cover (5km) and host species exhibit a strong positive association with prevalence of *P. knowlesi* in NHP (Table S15-17, Figure S15). Final adjusted OR for the full multivariable model can be visualised in Figure 3 (Table S14).

**Figure 3.**
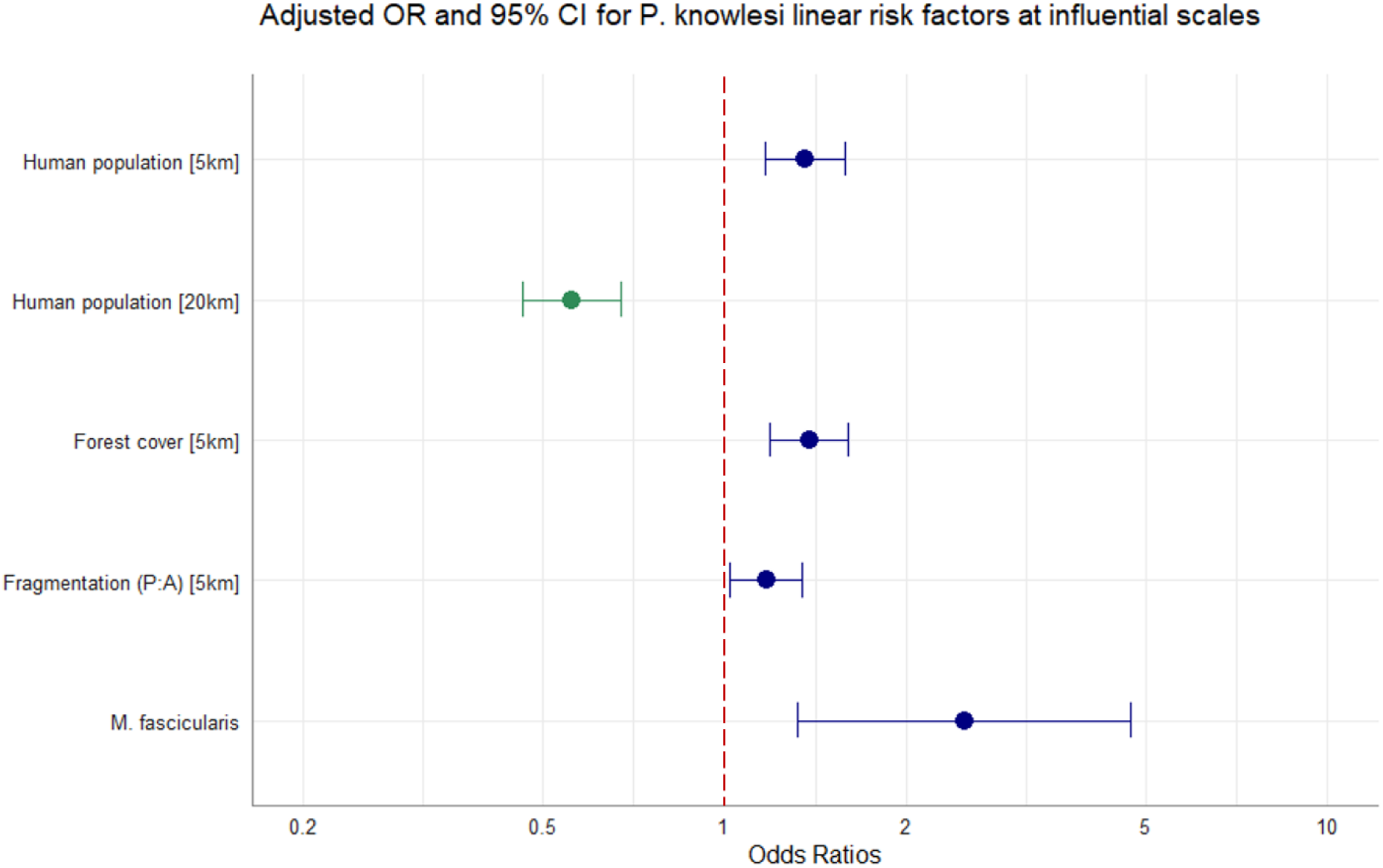
Multivariable regression results. Spatial scale denoted in square bracket. Canopy cover = %. Adjusted OR (dots) and CI95% (whiskers) for factors associated with P. knowlesi in NHPs at significant spatial scales. N=1354, accounting for replicate pseudo-sampling.

## DISCUSSION

Land use and land cover change is widely linked to spillover of zoonotic pathogens from sylvatic reservoirs into human populations, and pathogen prevalence in wildlife host species is key in driving the force of infection in spillover events. Our initial analyses found that for *Plasmodium knowlesi,* there is substantial spatial heterogeneity and prevalence in non-human primates varies markedly between regions of Southeast Asia (Zhang et al., 2016).

Consistent with our hypothesis that parasite density in primate hosts would be higher in areas experiencing habitat disturbance, we identified strong links between *P. knowlesi* in NHPs and measures of contemporaneous tree cover and habitat fragmentation. To our knowledge, this is the first systematic study to find evidence of landscape influencing the distribution of *P. knowlesi* prevalence in NHPs. Results offer evidence that *P. knowlesi* infection rates in NHPs are linked to changes in landscape across broad spatial scales, and that prevalence of *P. knowlesi* in reservoir species may be driving spillover risk across Southeast Asia. These findings could provide insight to improving surveillance of *P. knowlesi* and to the development of ecologically targeted interventions.

While previous studies have estimated that *P. knowlesi* infection would be chronic in all macaques, or as high as 50–90% for modelling *P. knowlesi* transmission in Malaysia (Brock et al., 2016), this data strongly suggests that this is not the case. Overall prevalence of *P. knowlesi* infection in all NHPs is markedly lower than usual estimates, emphasising the importance of accounting for absence data in estimations of prevalence. Considerable heterogeneity was identified between and within regional estimates for *P. knowlesi* across Southeast Asia, which likely reflects genuine differences according to distinct climates and habitats (Shearer et al., 2016). Malaysian Borneo was found to have an estimated prevalence over five-fold higher than West Malaysia. Crucially, such extreme prevalence estimates for NHPs in Borneo align with the known hotspot for human incidence of *P. knowlesi* (Cooper et al., 2020). By comparison, for Peninsular Malaysia, estimated prevalence is far lower than anticipated. Cases of human *P. knowlesi* do occur in West Malaysia, though transmission has been found to exhibit spatial clustering (Phang et al., 2020) which may correspond to pockets of high risk within the wider context of low prevalence of *P. knowlesi* in macaque populations. Regional trends in *P. knowlesi* also mask differences in infection rates between sample locations, driven by more localised factors.

Multiple studies reported finding *P. knowlesi* infections in wild macaques to be low or absent in peri-domestic or urbanised areas, attributed to the absence of vector species typically found in forest fringes (Brant et al., 2016; Chua et al., 2019; Manin et al., 2016). This pattern is seen in reports from Peninsular Malaysia (Saleh Huddin et al., 2019; Vythilingam et al., 2008), Singapore (Jeslyn et al., 2011; M. I. Li et al., 2021) and Thailand (Fungfuang et al., 2020; Putaporntip et al., 2010). The high heterogeneity of reports here suggests that the picture is even more complex. *P. knowlesi* infections may even vary between troops within a single study site, as was seen in the Philippines (Gamalo et al., 2019). Fine-scale interactions are unlikely to be captured by the scale of this study.

Ecological processes determining *P. knowlesi* infection are influenced by dynamic variables over multiple spatial scales (Cohen et al., 2016). We utilised a data-driven methodology to select variables at distances that capture maximum impact on *P. knowlesi* prevalence (Byrne et al., 2021; Fornace et al., 2019b), with tree cover and fragmentation influential at localised scales and human population density also exerting influence within wider radii. Contrary to previous studies on risk factors for human incidence of *P. knowlesi* (Fornace et al., 2019b, 2016), elevation was not found to be associated with *P. knowlesi* in NHPs at any scale. Vector and host species composition vary substantially across tropical ecotones, and it is likely that the study extent encompasses a range of putative vectors across different landscapes, such as those of the Minimus Complex in northern regions (Parker et al., 2015) or the recently incriminated *An.-collessi* and *An.-roperi* from the Umbrosus Group (de Ang et al., 2021). Given that the vector species driving sylvatic transmission remain elusive, it is conceivable that the elevation range covers multiple vector and host species niches and explains the lack of observed relationship between elevation and *P. knowlesi* in NHPs. Human population density was found to be significant at multiple distances, with contrasting effects on parasite prevalence in NHP. Previous studies have found a negative association between human density and vector density and biting rates in forested landscapes (Fornace et al., 2019a). Across wide spatial scales, increased vector density in less populated, more forested areas could generate higher parasite prevalence in NHPs. At the same time Long-tailed macaques, a species shown here to have higher prevalence rates, are notorious as nuisance animals and many of the available samples were collected opportunistically in urban areas, which might underly the observed positive association between localised high human density and higher prevalence in NHP. Whilst more data would be needed to understand this interaction, this further demonstrates the importance of using approaches to identify disease dynamics across multiple spatial scales (Brock et al., 2019).

A key finding is the link between high prevalence of *P. knowlesi* in primate host species with high degrees of habitat fragmentation. Habitat fragmentation is a key aspect of landscape modification, where large contiguous areas of habitat (for example, forests) are broken into a mosaic of smaller patches. This disturbs the ecological structure by increasing the density of fringes or ‘edges’, dynamic habitat often at the boundaries between natural ecosystems and human-modified landscapes (Borremans et al., 2019). Other studies have linked habitat fragmentation to increased generalist parasite density in primates. In Uganda, a higher prevalence and infection risk of protozoal parasites was observed in wild populations of red colobus primates (*Procolobus rufomitratus*) inhabiting fragmented forests compared to those in undisturbed habitat (Gillespie and Chapman, 2008). For *P. knowlesi,* creation of edge habitat is thought to favour vectors of the Leucosphyrus Complex (Davidson et al., 2019a; Hawkes et al., 2019). *Anopheles* spp. presence can be predicted by indices of fragmentation in Sabah, Borneo, with land cover changes creating more suitable micro-climate for larval habitats (Byrne et al., 2021), and an increased abundance of *An. balabacensis* found in forest fringes (Hawkes et al., 2019; Wong et al., 2015). Increasing landscape complexity results in increased density of edge habitat, with conceivably higher density of vectors in forest fringes. Therefore, preferential use of fringe habitat and high exposure to vectors in forest fringes may contribute to higher conspecific transmission of *P. knowlesi* between primates in increasingly fragmented habitats. This finding also lends clarity to landscape fragmentation as a risk factor for human exposure to *P. knowlesi* in Malaysian Borneo (Brock et al., 2019; Fornace et al., 2019b), with changes in relative host density, vector density and wildlife parasite prevalence in nascent forest fringes potentially enhancing the spillover of this disease system into human populations in fragmented habitats.

Conversely, we saw a strong association between high parasite prevalence and high tree canopy coverage. Given that a strong inverse relationship with fragmentation was observed, with high tree density correlating to low fragmentation indices and vice versa, this speaks to a trade-off between dense canopy cover and high habitat complexity and suggests an ‘ideal’ amount of habitat fragmentation that facilitates prevalence in primate hosts. For animals with larger home ranges, individual-based disease models combined with movement ecology approaches have shown that the most highly fragmented areas are less favourable for maintaining parasite transmission (White et al., 2018). In Sabah, individual macaques were shown to increase ranging behaviour in response to deforestation (Stark et al., 2019). Forest edge density also peaks at intermediate levels of land conversion (Borremans et al., 2019). With smaller habitat patches in maximally fragmented landscapes potentially insufficient to support macaque troops, this interplay between disease ecology and metapopulation theory may explain why both tree density and habitat fragmentation appear to pose a greater risk for simian *P. knowlesi.* Likewise, this may relate to the finding that in Borneo, larger forest patches (lower fragmentation indices) were associated with *P. knowlesi* spillover in Borneo (Fornace et al., 2019b). Overall, this finding offers an insight to mechanisms that underpin the increased force of infection of *P. knowlesi* that is associated with landscape change.

There are limitations to consider in the available data and interpretation of these findings. ‘Small-study effects’ were observed in the dataset, suggestive of a bias toward positive effect estimates (Stewart et al., 2012). This may be a result of data disaggregation and small studies creating artefactually higher estimates or may reflect true bias in data collection toward areas known to be endemic for *P. knowlesi* and convenience sampling of macaques. Assumptions have also been made that sample site equates to habitat, which may not reflect actual habitat use, and even accurate georeferenced data points are unlikely to entirely reflect surrounding habitat within the macaque home range. Variability in study designs and data reporting also impacted geospatial accuracy. Steps were taken to account for spatial bias by extracting covariates at randomly generated pseudo-sampling points. Whilst uncertainty cannot be eliminated, we demonstrate a robust methodology to accommodate for geographical uncertainty in ecological studies. Future investigations should prioritise systematic, georeferenced sampling across a range of landscape scenarios.

Results show important regional ecological trends, but broad geographic patterns may not be generalisable at individual levels, or to all putative host species in all geographic contexts (Zhang et al., 2016). Follow up studies should be conducted at higher spatial and temporal resolution to characterise the effect of local landscape configuration on wildlife *P. knowlesi* prevalence. Effects of fragmentation are likely to be dependent on land conversion type, species composition and surrounding matrix habitat (Fornace et al., 2019b). Use of perimeter: area ratio (PARA) as a fragmentation index was justified given high canopy coverage in study sites (Wang et al., 2014), though Edge Density (ED) or normalised Landscape Shape Index (nLSI) might be more appropriate in future analyses to account for variation in forest abundance. Specific land configurations have previously been linked to *P. knowlesi* exposure in Borneo (Fornace et al., 2019b), notably in areas where palm oil plantation is a dominant industry. Given this, broad forest classifications used here may mask important differences in *P. knowlesi* prevalence between land classes. As it was not possible to include contemporary land cover classifications in this analysis, future studies would also benefit from looking at specific habitat type (e.g., primary forest, agroforest, plantation).

## CONCLUDING REMARKS

Strong links have been identified between land use and land cover change and ecosystem perturbation that favours the transmission of vector-borne diseases (Loh et al., 2016). Prevalence of *P. knowlesi* in macaques is likely to be a crucial determinant of human infection risk, and more representative estimates of *P. knowlesi* prevalence derived here can better inform regional transmission risk models. This study also characterises landscape risk factors for heightened prevalence of *P. knowlesi* in NHPs. Findings provide evidence that *P. knowlesi* in primate hosts is partly driven by landscape modification across Southeast Asia. While the full complexity is not captured by the covariates used, it is clear that *P. knowlesi* infection in NHPs is not restricted to densely forested areas. This study also demonstrates the utility of systematic meta-analysis tools and remote-sensing datasets in the investigation of macroecological disease trends, in conjunction with methods to standardise a spatially heterogeneous dataset and data-driven selection of spatial scales. Gaps identified in data reporting should inform more systematic and localised primatological surveys to disentangle precise mechanisms. Notwithstanding limitations, this study highlights the marked spatial heterogeneity and role of landscape complexity in driving *P. knowlesi* infection rates in NHPs. Given the clear intersection between human epidemiology and wildlife ecology, it is essential that infection dynamics within wildlife reservoirs are considered in future public health interventions.

## METHODS

### Study site

This study focused on the simian malaria *Plasmodium knowlesi* across Southeast Asia, within 28°30’00.0″N, 92°12’00.0″E and 11°00’00.0″S, 141°00’00.0″E. Climate mainly corresponds to the equatorial tropical zone, with high temperatures and high humidity.

### Data assembly

A systematic literature review was conducted under the CoCoPop framework (Condition, Context, Population) (Cuenca et al., 2022; Munn et al., 2015). All studies identified in the literature review were screened for data on NHPs with a confirmed *P. knowlesi* diagnosis or absence data (zero counts of *P. knowlesi* with appropriate diagnostic methods). Exclusion criteria included (a) studies exclusively relying on microscopy (Antinori et al., 2013) (b) laboratory, animal model or experimental infection studies (c) data from outside of Southeast Asia. No limit was set on the temporal range for primate survey records. Duplicate records reporting results from the same surveys were removed, with one record per survey retained. Critical appraisal of the studies was conducted using the Joanna Briggs Institute (JBI) checklist for prevalence studies (Munn et al., 2015) (see Supplementary Information (SI) for details and criteria). A flowchart of the selection process is illustrated in Figure S3, with a full list of articles included provided in Table S2.

Primary outcome was defined as *P. knowlesi* prevalence (*p,* proportion positive for *P. knowlesi* infection from *n* sampled NHPs). For each independent primate study, the following variables were extracted: year of data collection, primate species sampled, primate status (wild/captive), diagnostic test (PCR/sequencing) and target gene(s), sampling method (routine/purposive), number of *P. knowlesi* positive samples, number of *Plasmodium* spp. positive samples, total number of primates tested and geographical information.

In most studies identified, study site was only geolocated to a geographic area or descriptive location. Geolocation was assigned at the lowest available level of administrative polygon (i.e., district/state/country) by cross-referencing reported sampling location with GADM (v3.6) administrative boundaries. If specific location was given, GPS coordinates were assigned via Google Maps. For data visualisation, point coordinates were plotted in QGIS (3.10.14) and R (4.1.0) software.

### Meta-analysis of *P. knowlesi* prevalence

Meta-analysis was conducted using methods that are standard in the analysis of human disease prevalence for individual participant datasets (IDP) (Liberati et al., 2009; Stewart et al., 2012). Data were disaggregated by geographic location (site) and primate species, to illustrate variance in prevalence by survey unit (Stewart et al., 2012). One-stage meta-analysis is considered appropriate for studies where the outcome may be infrequent, so data was included in a single model under the ‘DerSimonian and Laird’ variance estimator (Munn et al., 2015). Sensitivity analyses were conducted to compare methods for the back-transformation of prevalence estimates. For studies where prevalence estimates tend towards 0% or 100%, variance tends towards 0. To stabilise the variance and enable back-transformation of zero prevalence records, logit method was selected for the transformation of prevalence, with the inverse variance method used for individual study weights (see SI for details).

Overall heterogeneity of prevalence records was assessed using the I^2^ statistic (Hippel, 2015), a relative estimate of true between-study variance. Sub-group analysis was conducted according to geographic region, with the heterogeneity of reported prevalence within regional sub-groups assessed using prediction intervals derived from the τ ^2^ statistic. Small-study effects, including selection and publication biases, were assessed by examining funnel plots and imputing ‘missing’ estimates using the trim-and-fill method (Lin and Chu, 2018). Full rationale and details of small-study effect assessments can be found in Supplementary Information.

### Remote sensing data

Satellite-derived remote sensing datasets were used to assemble local environmental and anthropogenic covariates. Gridded UN-adjusted human population estimates were assembled at 1km resolution from WorldPop (WorldPop, 2018). Elevation data was obtained from NASA SRTM 90m Digital Elevation Database v4.1 (CGIAR-CSI) (Jarvis, A., H.I. Reuter, A. Nelson, 2008) with a spatial resolution of 1km. Contemporaneous tree cover was derived from Hanson’s Global Forest Watch (30m) (Hansen et al., 2013), extracted for every year between 2006–2020.Tree cover was classified as ≥50% crown density, and then matched to primate data by sample site geolocation and by year of sample collection to account for rapid forest loss (SI, Figure S7). Where a broad timeframe of sampling was provided (≥3 years), median year was used. Full details for variable selection and processing can be found in Supplementary Information (Table S11–12, Figure S9).

Perimeter: area ratio (PARA, ratio of patch perimeter length to patch surface area) of given land class is a key metric for habitat conversion, where a higher PARA provides a measure of boundary complexity and indicates a more fragmented landscape (McGarigal and Cushman, 2012). Mean PARA was extracted from canopy cover within circular buffers. Habitat fragmentation has been shown to correlate with disease transmission parameters (Borremans et al., 2019; Faust et al., 2018), but definitions often lack precision and can be considered with respect to ‘separation effects’ (division and isolation of patches) and ‘geometric effects’ (changes to ratios of perimeter and core habitat) (Wilkinson et al., 2018). PARA provides a measure of edge density within the buffer area (PARA>0) and has been shown to provide a good index of fragmentation and good discrimination of spatial aggregation across areas where habitat abundance (tree canopy cover) is high. (Wang et al., 2014) (SI, Table S12, Figure S10).

### Covariate assembly

For studies with exact GPS coordinates, precise environmental data at a single site could be obtained. For surveys published without GPS coordinates, there is considerable geographic uncertainty in the exact sampling location (SI, Table S7). Uncertainty in the spatial and environmental determinants of prevalence generates a sampling bias, with the precision of covariates correlated to certain studies. Use of a single centroid proxy site is standard procedure, but often generates erroneous estimates in large or heterogenous sampling units(Cheng et al., 2021). Alternative strategies were employed to account for and mitigate the effect of spatial uncertainty and spatial bias. Each prevalence observation was replicated and assigned a random sample of environmental realisations. 10 random sampling points were generated within the sampling area provided by the study, and covariates were extracted at each proxy sampling site (SI, Table S8). Selection of random points was validated by visual inspection of the stability of model coefficients with the inclusion of an increasing number of points. Number of points was selected conservatively at the point where coefficients stabilised (n=10).

For every georeferenced sampling point, mean values for all selected covariates were extracted within buffer radii at 5km, 10km and 20km (SI, Figure S11). Buffer area sizes were selected to investigate multiple spatial scales over which associations between risk factors and *P. knowlesi* prevalence might occur. A minimum radius of 5km was chosen to approximate the maximum ranging distance for *M. fascicularis* (Waxman et al., 2014), with wider radii (10–20km) included to account for the geographic uncertainties in areal data. Flowchart of data processing chain can be found in Supplementary Information (Figure S8).

### Analysis of environmental risk factors

Generalised linear mixed-effect regression models (GLMM) were fitted to NHP prevalence data using a binomial distribution with a logit link. To account for within-study correlation in reported average prevalence, a unique identifier combining author and study was included as a random intercept in all models. Artificial inflation of sample size in the replicated data (10 pseudo-sampling sites for data geolocated to administrative areas) was accommodated by reducing individual observation weights to 1/10^th^ within the model.

Each covariate at each spatial scale was assessed for inclusion in the multivariable model based on bivariable analysis and a criterion of *p* >0.2 under likelihood ratio tests (LRT) (Table S13). A quadratic term for the fragmentation index ‘PARA’ was included to account for possible nonlinearity. Multicollinearity among independent predictors at multiple scales was examined via variance inflation factors (VIF). The VIF of each predictor variable was examined following a stepwise procedure, starting with a saturated model and sequentially excluding the variable with the highest VIF score from the model. Stepwise selection continued in this manner until the entire subset of explanatory variables in the global model satisfied a moderately conservative threshold of VIF ≤6 (Rogerson, 2001). Qualifying variables obtained were then assessed for model inclusion using a backward stepwise strategy, removing variables with the highest *p* value (LRT) until a pre-defined threshold of α <0.05. Spearman’s rank tests were conducted on the selected variables to observe residual correlation, plotted as a correlation matrix (Figure S14).

Fully adjusted OR for associations between environmental covariates and *P. knowlesi* prevalence were derived from the final multivariable GLMM with *p* values derived from LRT. Spatial sensitivity analyses were conducted by excluding data points from administrative boundaries outside a reasonable size or above a reasonable threshold of environmental certainty, according to the standard deviation (SD) of the covariate values within each set of 10 environmental realisations (Table S15 and Table S16). Ecological sensitivity analyses were conducted by removing data points that fall outside areas with high predicted probability of occurrence for *Macaca fascicularis, Macaca nemestrina* and *Macaca leonina* and running regression analyses on the constrained dataset (Moyes et al., 2016) (Table S17, Figure S15).

### Ethics

Ethics approval was not required for this research, following assessment by the LSHTM Research Governance & Integrity Office (LSHTM MSc Ethics Ref: 25429).

## ACKNOWLEDGEMENTS

Research was supported by the Sir Henry Dale Fellowship, jointly funded by the Wellcome Trust and the Royal Society (Grant Number 221963/Z/20/Z). Data collected in Peninsular Malaysia by the Faculty of Veterinary Medicine, Universiti Putra Malaysia, was funded by the Ministry of Higher Education Malaysia (Grant LRGS/1/2018/UM/01/1/). Additional funding was supported by the World Health Organization.

