## Supplementary Information for "Landscape drives zoonotic malaria prevalence in non-human primates"

#### APPENDIX A. Data assembly

Prior to conducting the study, a review of current literature was constructed to find articles related to “Plasmodium knowlesi” or to both “malaria” and “primate”, including synonyms and sub-headings. The search was elaborated to specify environmental factors (Figure S1). The following databases were searched:

- Medline
- Embase
- Web of Science

Provisional data were extracted using a standardised form (Figure S2) using standardised definitions (Table S1), from which an initial set of studies were identified for this investigation. Duplicate records (confirmed/suspected to be the same specimens) were removed, with one record retained (Figure S3)

Embase Classic+Embase <1947 to 2021 July 02>

| | | |
|---|---|---|
| 1 | (monkey* or macaca* or macaque* or primate* or zoono* or simian).mp. | 284859 |
| 2 | Zoonosis/ep, et [Epidemiology, Etiology] | 3000 |
| 3 | Catarrhini/ or Cercopithecidae/ or Cercopithecinae/ or Macaca/ | 29691 |
| 4 | 1 or 2 or 3 | 285484 |
| 5 | Malaria/ep [Epidemiology] | 11117 |
| 6 | (malaria or plasmodium or inui or cynomolgi or coatneyi or knowlesi).mp. | 148966 |
| 7 | 5 or 6 | 148966 |
| 8 | 4 and 7 | 4577 |
| 9 | Plasmodium knowlesi/ | 1349 |
| 10 | plasmodium knowlesi.mp. | 1734 |
| 11 | 9 or 10 | 1734 |
| 12 | 8 or 11 | 5439 |
| 13 | (forest* or biodivers* or ecolog* or fragment* or deforest* or anthropogenic or environment* or climat*).mp. | 2514624 |
| 14 | 12 and 13 | 710 |

***Figure S1.*** *Search strategy for background research*

| Species | Presence | # | Date | Country | Location | Latitude | Longitude | Source ID |
|---|---|---|---|---|---|---|---|---|

***Figure S2.*** *WHO Report primate data extraction form*

***Table S1.*** *Standardised definitions for qualitative primate characteristics*

| Variable | Category | Definition |
|---|---|---|
| Sampling | Routine | Animals collected for surveillance purposes or extracted from human–conflict zones; data collected opportunistically [1] |
| | Study | Animals captured and sampled specifically for a study of Plasmodium spp and/or P. knowlesi |
| Status | Captive | Animal resident in sanctuary or conservation park |
| | Wild | Free–living animal, not registered/resident in any sanctuary |
| | Sanctuary | A wildlife sanctuary/rehabilitation centre housing key primate species [2] |
| Area | Forest | Areas that are uninhabited with extensive tree cover |
| | Peri–domestic | As defined by the author. Example definitions as follows:<br>· Rural areas (areas with low human density, close to secondary/scrub forest) [3]<br>· Public nature reserve park [4]<br>· 2 km from longhouse communities [5]<br>· Wild Long-tailed macaque samples collected based on their proximity to humans [6] |
| | Agricultural | Animal located in agricultural areas, predominantly mono culture (e.g. orchard, plantation) [3] |
| | Urban | As defined by the author. Generally, areas with high human population density [7] |

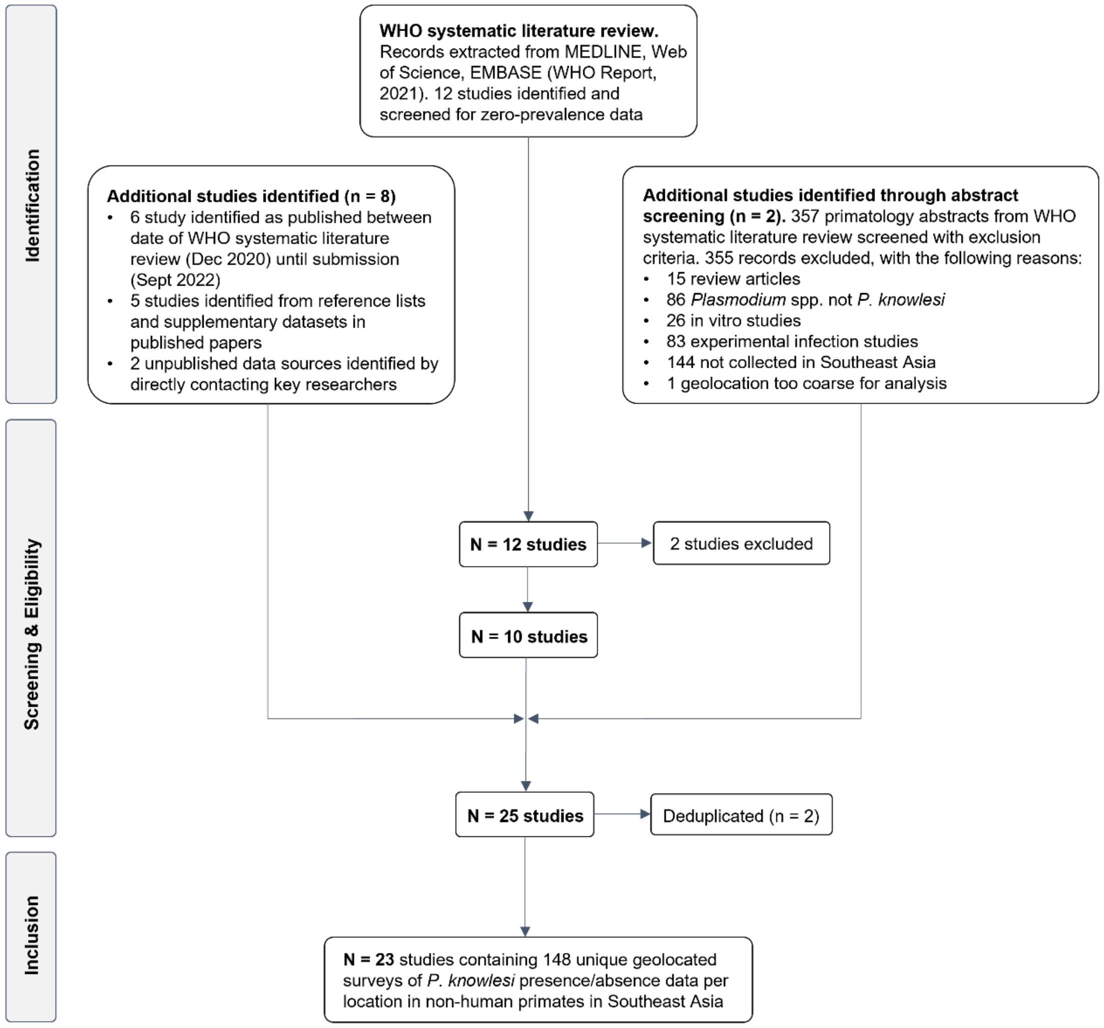

***Figure S3.*** *Flow chart illustrating study selection process*

***Table S2.*** *Characteristics of the included studies*

| Author | Year(s) | Country/region | N* | Sample† | Diagnostic | Target gene(s) | Primer |
|---|---|---|---|---|---|---|---|
| Lee et al., 2011 | 2004–2008 | Malaysia/Borneo | 108 | Study | Nested PCR | *SSU rRNA/csp/mtDNA* | Kn1f/Kn3r |
| Seethamchai et al., 2008 | 2006 | Thailand | 99 | Study | Sequencing | *A type SSU rRNA/cytb* | • |
| Vythilingam et al., 2008 | 2007 | Malaysia/Peninsular | 145 | Study | PCR/Sequencing | *SSU rRNA/csp* | Pmk8/Pmkr9 |
| Zhang et al., 2016 | 2007 | Singapore | 40 | Study | PCR | • | • |
| | 2007–2010 | Indonesia/Sumatra | 70 | Study | PCR | • | • |
| | 2011 | Cambodia | 54 | Study | PCR | • | • |
| | 2012 | Philippines | 68 | Study | PCR | • | • |
| | 2015 | Laos | 44 | Study | PCR/Sequencing | *SSU rRNA* | PK18SF/PK18SRc |
| Jeslyn et al., 2011 | 2008 | Singapore | 13 | Routine | PCR/Sequencing | *SSU rRNA /csp* | Pmk8/Pmkr9 |
| Ho et al., 2010 | 2008 | Malaysia/Peninsular | 107 | Routine | Nested PCR | *SSU rRNA* | Pmk8/Pmkr9 |
| Li et al., 2021 | 2008–2017 | Singapore | 1039 | Routine | Nested PCR | *SSU rRNA* | Pmk8/Pmkr9 |
| Putaporntip et al., 2010 | 2009 | Thailand | 655 | Study | Sequencing | *cytb* | • |
| Chang et al., 2011 | 2010 | Myanmar | 45 | Study | PCR | *SSU rRNA* | • |
| Muehlenbein et al., 2015 | 2010 | Malaysia/Borneo | 41 | Study | PCR | *mtDNA/AMA-1/MSP-1* | • |
| Zarith et al., 2021‡ | 2010–2017 | Malaysia/Peninsular | 1587 | Routine | Nested PCR | *SSU rRNA* | • |
| Unpublished, 2013†† | 2013 | Malaysia/Peninsular | 15 | Study | PCR | • | • |
| Unpublished, 2015†† | 2013–2016 | Malaysia/Borneo | 25 | Study | Nested PCR | *cytb* | PKCBF/PKCBR |
| Saleh Huddin et al., 2019 | 2014 | Malaysia/Peninsular | 415 | Study | PCR/Sequencing | *SSU rRNA* | Pmk8/Pmkr9 |
| Akter et al., 2015 | 2015 | Malaysia/Peninsular | 70 | Routine | PCR/Sequencing | *A-type SSU rRNA* | Pmk8/Pmkr9 |
| Amir et al., 2020 | 2016 | Malaysia/Peninsular | 103 | Routine | Nested PCR | *SSU rRNA* | PkF1140/PkR1550 |
| Gamalo et al., 2019 | 2017 | Philippines | 95 | Study | Nested PCR | *SSU rRNA* | Kn1f/Kn3r |
| Fungfuang et al., 2020 | 2017–2019 | Thailand | 93 | Study | Nested PCR | *SSU rRNA* | Kn1f/Kn3r |
| Nada-Raja et al., 2022 | 2018 | Malaysia/Borneo | 73 | Study | Nested PCR | *SSU rRNA/csp/mtDNA* | Kn1f/Kn3r |
| Yusuf et al., 2022 | 2016-2019 | Malaysia | 419 | Study | Nested PCR | *SSU rRNA* | Kn1f/Kn3r |
| Zamzuri et al., 2022 | 2018 | Malaysia/Peninsular | 212 | Routine | PCR | • | • |
| Kaewchot et al., 2022 | 2019 | Thailand | 649 | Study | Nested PCR | *SSU rRNA* | Pmk8/Pmkr9 |
| Salwati et al., 2017 | 2015 | Indonesia/Sumatra | 38 | Study | PCR/Sequencing | • | • |

*N=number of primates sampled
†Animal trapped either on routine or study basis
‡Unpublished, personal correspondence (p/c)
††Danau Girang Field Centre, p/c from Dr Salgado Lynn

Of the 87 records reporting presence of *P. knowlesi,* only 22 records (containing 248 *P. knowlesi* positive macaques) report whether *P. knowlesi* infection was a mono-infection or mixed infection with other simian *Plasmodium* spp. With a low proportion of data represented, this was deemed insufficient to conduct any further investigations.

*Macaca fascicularis* is the predominant species tested. However, reports also include *M. nemestrina* (6.1%, n=9/148; 527 macaques)[5,8–10], *M. arctoides* (1.4%, n=2/148; 36 macaques)[10,11], *M. leonina* (n=1/148; 25 macaques)[11]*, Trachypithecus obscurus* (Dusky leaf monkey) (n=1/148; 7 tested) and unspecified species from the *Presbytis* genus (n=1/148; 2 tested) (Table S6). One study additionally sampled 1 *Presbytis melalophos* (Black-crested Sumatran langur)[12], but species-specific *P. knowlesi* was not reported (Table S3, S4).

***Table S3.*** *Published studies of* P. knowlesi *infections (n) in non-human primate species collected (N) in Southeast Asia, grouped by region and author*

| Region | Species | | | | Total | Ref. |
|---|---|---|---|---|---|---|
| | *M. fascicularis* | *M. nemestrina* | *M. arctoides* | Other | | |
| Peninsular Malaysia | 25 /107 | • | • | • | | 13 |
| | 48 /415 | • | • | • | | 6 |
| | 21 /70 | • | • | • | | 14 |
| | 11 /98 | 0 /5 | • | • | | 8 |
| | 0 /15 | • | • | • | 473 /3069 | 15 |
| | 10 /145 | • | • | • | | 12 |
| | 215 /1587 | • | • | • | | 3 |
| | 66 /415 | • | • | • | | YF |
| | 74 /207 | 3 /5 | • | • | | ZZ |
| Borneo | 4 /26 | 2 /15 | • | • | | 9 |
| | 71 /82 | 13 /26 | • | • | | 5 |
| | 18 /25 | • | • | • | 119 /251 | 16 |
| | 7 /45 | 2 /28 | • | • | | NR |
| | 2 /4 | • | • | • | | YF |
| Sumatra | 0 /70 | • | • | • | 6 /108 | 17 |
| | 5 /32 | 1 /4 | • | 0 /2† | | SW |
| Thailand | 1 /195 | 5 /449 | 0 /4 | 1 /7‡ | | 10 |
| | 0 /21 | • | • | • | | 18 |
| | 0 /4 | • | • | • | 8 /1496 | 11 |
| | 0 /32 | 0 /25* | 1 /32 | • | | 11 |
| | 0 /78 | • | • | • | | 18 |
| | 0 /649 | • | • | • | | KT |
| Philippines | 18 /95 | • | • | • | 18 /163 | 2 |
| | 0 /68 | • | • | • | | 17 |
| Singapore | 3 /13 | • | • | • | | 4 |
| | 145 /1039 | • | • | • | 148 /1092 | 7 |
| | 0 /40 | • | • | • | | 17 |
| Laos | 1 /44 | • | • | • | 1 /44 | 17 |
| Cambodia | 0 /54 | • | • | • | 0 /54 | 17 |
| Myanmar | 0 /45 | • | • | • | 0 /45 | 19 |
| Total | 743 /5720 | 26 /557 | 1 /36 | 1 /9 | **773 /6322** | |

**Macaca leonina* (Northern Pig-tailed macaque, recently classified as separate species)
†*Presbytis* spp.
‡*Trachypithecus obscurus* (Dusky leaf monkey)

***Table S4.*** *Characteristics of primates tested and number/percentage of confirmed* P. knowlesi *infections (Pk+)*

| | | N | (%)* | *Pk+* | *Pk+* (%)† | CI95%‡ |
|---|---|---|---|---|---|---|
| Species | *M. fascicularis* | 5720 | (90.5%) | 745 | 13.0% | (12.2–13.9) |
| | *M. nemestrina* | 532 | (8.4%) | 26 | 4.9% | (3.4–7.1) |
| | *M. leonina* | 25 | (0.4%) | 0 | 0.0% | (0.0–13.3) |
| | *M. arctoides* | 36 | (0.6%) | 1 | 2.8% | (0.5–14.2) |
| | *T. obscurus* | 7 | (0.1%) | 1 | 14.3% | (2.6–51.3) |
| | *Presbytis* spp. | 2 | (0.03%) | 0 | 0.0% | (0.0–65.8) |
| Area | Forest | 1740 | (27.5%) | 253 | 14.5% | (13.0–16.3) |
| | Agriculture | 491 | (7.8%) | 72 | 14.7% | (11.8–18.1) |
| | Peri-domestic | 2192 | (34.7%) | 341 | 15.6% | (14.1–17.1) |
| | Urban | 1143 | (18.1%) | 56 | 4.9% | (3.8–6.3) |
| | Sanctuary | 109 | (1.7%) | 5 | 4.6% | (2.0–10.3) |
| | Unspecified | 647 | (10.2%) | 46 | 7.1% | (5.4–9.4) |
| Status | Wild | 6183 | (97.8%) | 768 | 12.4% | (11.6–13.3) |
| | Captive | 139 | (2.2 %) | 5 | 3.6% | (1.5–8.1) |
| Region | Pen. Malaysia | 3069 | (48.5%) | 473 | 15.4% | (14.2–16.7) |
| | Borneo | 251 | (4.0%) | 119 | 47.4% | (41.3–53.6) |
| | Sumatra | 108 | (1.7%) | 6 | 5.5% | (2.6–11.6) |
| | Thailand | 1496 | (23.7%) | 8 | 0.5% | (0.3–1.1) |
| | Philippines | 163 | (2.6 %) | 18 | 11.0% | (7.1–16.8) |
| | Singapore | 1092 | (17.3%) | 148 | 13.6% | (11.7–15.7) |
| | Cambodia | 54 | (0.9%) | 0 | 0.0% | (0.0–6.6) |
| | Laos | 44 | (0.7%) | 1 | 2.3% | (0.4–11.8) |
| | Myanmar | 45 | (0.7%) | 0 | 0.0% | (0.0–7.9) |
| Total | | 6322 | (100%) | 773 | 12.2% | (11.4–13.1) |

*Percentage of total number of primates tested (column %)

†Proportion of N positive for *P. knowlesi* (row %)

‡95% confidence interval (CI95%) calculated in R using count and sample size (binomial distribution)

### APPENDIX B. Quality appraisal

Quality was assessed using the Joanna Briggs Institute (JBI) Critical Appraisal tool for prevalence studies[20]. Studies were assessed on nine standardised criteria used to inform inclusion in the meta-analysis. Full criteria and examples of scoring are given in Tables S5 and S6. Studies assessed to be of lower quality were those that omitted key information about the sampling method[15]. Two studies were considered to be of higher quality owing to robust sampling and completeness of evidence[5,6]. Given the objective to assess variation in reported prevalence, and in considering the limited usefulness of criteria designed for human participants, the reliable diagnostic methods identified in all studies and the appreciable limitations in surveying wild animals, data from all studies (n=148 estimates) were included for further analyses.

***Table S5**. JBI criteria for assessing bias in meta-analyses of prevalence studies*

| Criteria | | Yes | No | Unclear | N/A |
|---|---|---|---|---|---|
| Q1 | Was the sample frame appropriate to address the target population? | | | | |
| Q2 | Were study participants sampled in an appropriate way? | | | | |
| Q3 | Was the sample size adequate? | | | | |
| Q4 | Were the study subjects and the setting described in detail? | | | | |
| Q5 | Was the data analysis conducted with sufficient coverage of the identified sample? | | | | |
| Q6 | Were valid methods used for the identification of the condition? | | | | |
| Q7 | Was the condition measured in a standard, reliable way for all participants? | | | | |
| Q8 | Was there appropriate statistical analysis? | | | | |
| Q9 | Was the response rate adequate, and if not, was the low response rate managed appropriately? | | | | |

***Table S6**. Example rationale for quality appraisal*

| Sample question | | Example | Assessment |
|---|---|---|---|
| Q1 | Was the sample frame appropriate to address the target population? | Wild animal<br>Captive animal<br>Not specified | Yes<br>No<br>Uncertain |
| Q2 | Were study participants sampled in an appropriate way? | Trapped for study<br>Routine collection<br>Not specified | Yes<br>No<br>Uncertain |
| Q9 | Was the response rate adequate? | Primate data | N/A |

### APPENDIX C. Meta analysis

#### *D.1. Subgroup analysis*

Sub-group analysis by region was conducted under a random-effects model. Pooled estimates are then back-transformed for interpretation. The Freeman-Tukey double arcsine method is recommended in the transformation of prevalence[20]. However, recent studies have found that back-transformation of the Freeman-Turkey method can generate misleading results, owing to the requirement for a global sample size for inversion[21]. A sensitivity analysis conducted using the logit transformation and untransformed proportions (Table S9) revealed a deficit in the back-transformation of the pooled prevalence estimate for Thailand under the Freeman-Tukey transformation, generating a null point estimate. To avoid this error, meta-analysis was conducted under the logit transformation with the inverse variance estimator to account for individual study weighting.

***Table S9.*** *Sensitivity analysis for transformation of P. knowlesi prevalence estimate under random-effects model, shown overall and for Thailand subgroup analysis*

| Method | Overall (*k*=148) | | Subgroup (Thailand, *k*=21) | |
|---|---|---|---|---|
| | P* | CI95% | P | CI95% |
| Freeman-Turkey double arcsine | 0.0943 | (0.0641–0.1284) | **0.0000** | **(0.0000–0.0000)** |
| Logit | 0.1199 | (0.0935–0.1526) | 0.0199 | (0.0113–0.0346) |
| Untransformed | 0.1415 | (0.1101–0.1730) | 0.0022 | (0.0000–0.0059) |

*D.3. Small study effects*

Small study effects, including selection and publication bias, were assessed by examining funnel plots and imputing ‘missing’ estimates using the trim-and-fill method[22]. Funnel plots were generated using both SE and study size as metrics of variance, as study size has been shown to be more accurate for meta-analyses of proportions where raw estimates tend towards 0 or 1[20]. Funnel plots for the disaggregated dataset are shown using SE as the variance estimate in Figure S4. Asymmetry is highlighted by the trim-and-fill interpolation method, which provides an estimate of missing data and added an additional 54 imputed points to the plot.

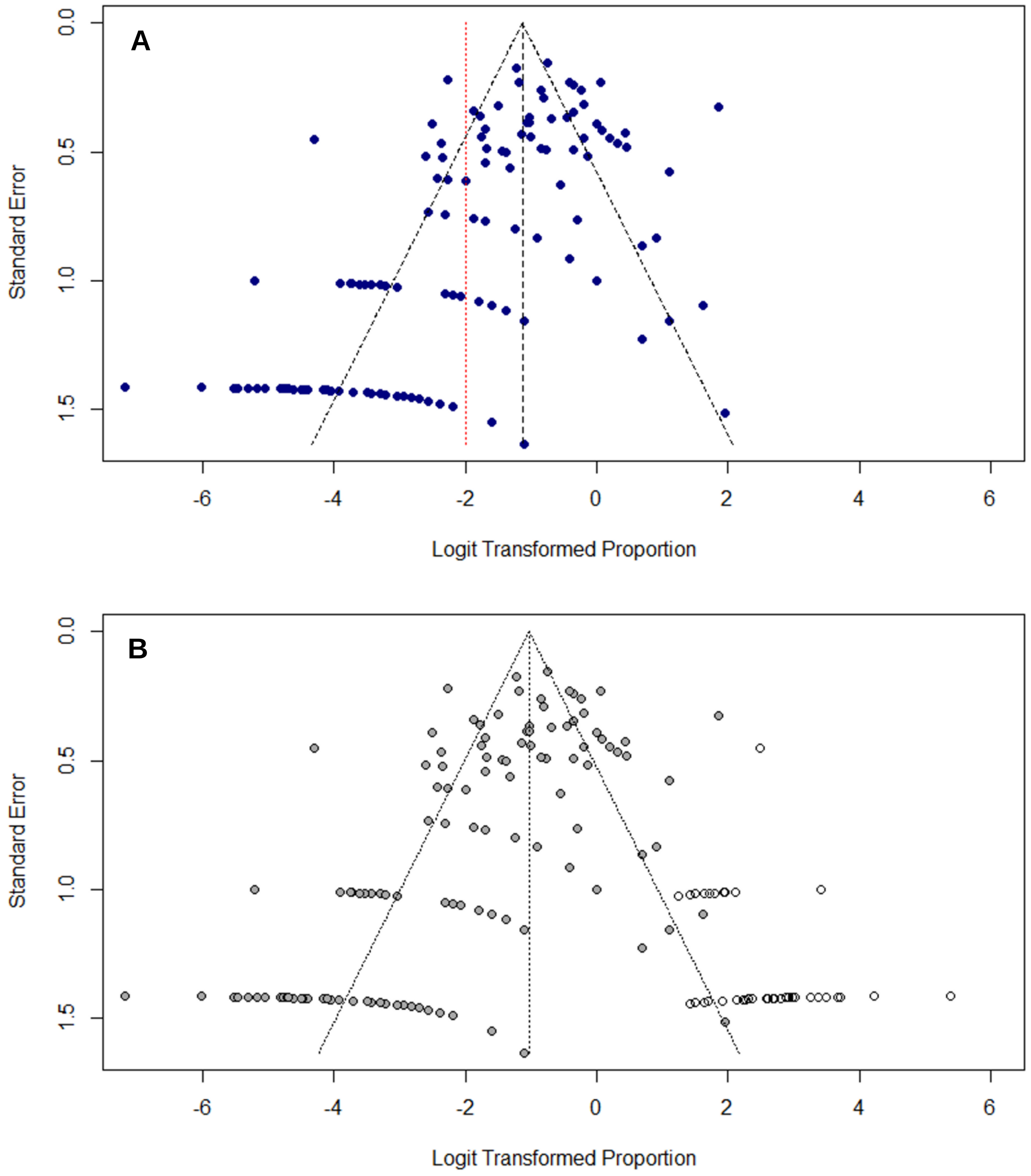

***Figure S4.*** *(A) Funnel plot of transformed prevalence (%) against standard error (SE) for study sites (B) Funnel plot with imputed data to illustrate asymmetry using trim-and-fill method*

*D.4. Regional prevalence estimates*

Meta-analysis was conducted with data disaggregated by survey location and primate species ($k$=148). Forest plot of individual study prevalence, presented with pooled regional prevalence estimates and relative sampling effort for the disaggregated data can be visualised in Figure S7.

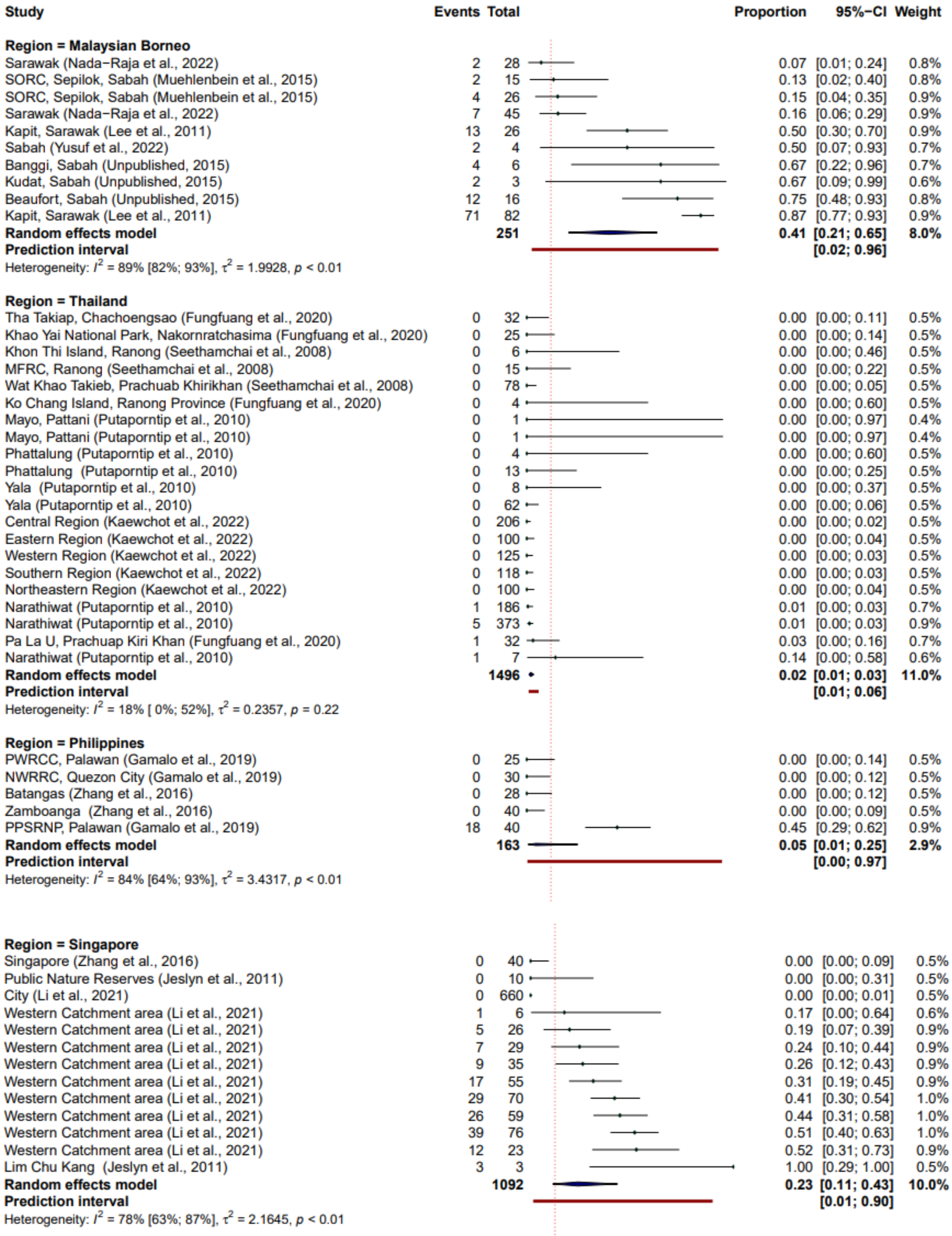

*Figure S6 (cont.)*

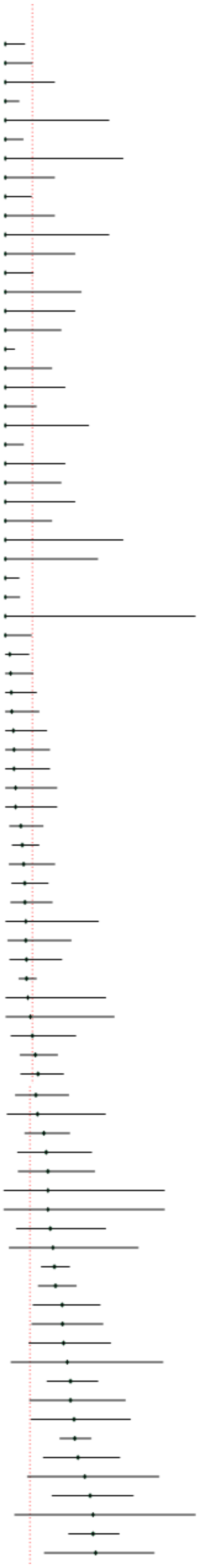

| Study | Events | Total | Proportion | 95% CI | Weight |
|---|---|---|---|---|---|
| **Region = Peninsular Malaysia** | | | | | |
| Selangor (Vythilingam et al., 2008) | 0 | 41 | 0.00 | [0.00; 0.09] | 0.5% |
| Kuala Lumpur (Vythilingam et al., 2008) | 0 | 29 | 0.00 | [0.00; 0.12] | 0.5% |
| Kuala Kubu Bharu, Hulu Selangor (Unpublished, 2013) | 0 | 15 | 0.00 | [0.00; 0.22] | 0.5% |
| Kedah (Saleh Huddin et al., 2019) | 0 | 59 | 0.00 | [0.00; 0.06] | 0.5% |
| Padang Besar (Zarith et al., 2022) | 0 | 6 | 0.00 | [0.00; 0.46] | 0.5% |
| Arau (Zarith et al., 2022) | 0 | 45 | 0.00 | [0.00; 0.08] | 0.5% |
| Gelugor (Zarith et al., 2022) | 0 | 5 | 0.00 | [0.00; 0.52] | 0.5% |
| Hulu Selangor (Zarith et al., 2022) | 0 | 15 | 0.00 | [0.00; 0.22] | 0.5% |
| Kuala Langat (Zarith et al., 2022) | 0 | 30 | 0.00 | [0.00; 0.12] | 0.5% |
| Sepang (Zarith et al., 2022) | 0 | 15 | 0.00 | [0.00; 0.22] | 0.5% |
| Besut (Zarith et al., 2022) | 0 | 6 | 0.00 | [0.00; 0.46] | 0.5% |
| Hulu Perak (Zarith et al., 2022) | 0 | 10 | 0.00 | [0.00; 0.31] | 0.5% |
| Kubang Pasu (Zarith et al., 2022) | 0 | 28 | 0.00 | [0.00; 0.12] | 0.5% |
| Bachok (Zarith et al., 2022) | 0 | 9 | 0.00 | [0.00; 0.34] | 0.5% |
| Kuala Krai (Zarith et al., 2022) | 0 | 10 | 0.00 | [0.00; 0.31] | 0.5% |
| Pasir Puteh (Zarith et al., 2022) | 0 | 13 | 0.00 | [0.00; 0.25] | 0.5% |
| Klang Valley (S/KL) (Saleh Huddin et al., 2019) | 0 | 88 | 0.00 | [0.00; 0.04] | 0.5% |
| Larut (Zarith et al., 2022) | 0 | 16 | 0.00 | [0.00; 0.21] | 0.5% |
| Bagan Datuk (Zarith et al., 2022) | 0 | 12 | 0.00 | [0.00; 0.26] | 0.5% |
| Kuantan (Zarith et al., 2022) | 0 | 25 | 0.00 | [0.00; 0.14] | 0.5% |
| Pekan (Zarith et al., 2022) | 0 | 8 | 0.00 | [0.00; 0.37] | 0.5% |
| Muallim (Zarith et al., 2022) | 0 | 44 | 0.00 | [0.00; 0.08] | 0.5% |
| Jelebu (Zarith et al., 2022) | 0 | 12 | 0.00 | [0.00; 0.26] | 0.5% |
| Hilir Perak (Zarith et al., 2022) | 0 | 13 | 0.00 | [0.00; 0.25] | 0.5% |
| Bukit Kayu Hitam (Zarith et al., 2022) | 0 | 10 | 0.00 | [0.00; 0.31] | 0.5% |
| Melaka Tengah (Zarith et al., 2022) | 0 | 16 | 0.00 | [0.00; 0.21] | 0.5% |
| Temerloh, Pahang (Amir et al., 2020) | 0 | 5 | 0.00 | [0.00; 0.52] | 0.5% |
| Port Dickson (Zamzuri et al., 2020) | 0 | 7 | 0.00 | [0.00; 0.41] | 0.5% |
| Kedah (Yusuf et al., 2022) | 0 | 59 | 0.00 | [0.00; 0.06] | 0.5% |
| Selangor (Yusuf et al., 2022) | 0 | 56 | 0.00 | [0.00; 0.06] | 0.5% |
| Putrajaya (Yusuf et al., 2022) | 0 | 2 | 0.00 | [0.00; 0.84] | 0.5% |
| Kuala Lumpur (Yusuf et al., 2022) | 0 | 30 | 0.00 | [0.00; 0.12] | 0.5% |
| Machang (Zarith et al., 2022) | 1 | 51 | 0.02 | [0.00; 0.10] | 0.7% |
| Klang (Zarith et al., 2022) | 1 | 43 | 0.02 | [0.00; 0.12] | 0.7% |
| Batu Pahat, Johor (Amir et al., 2020) | 1 | 38 | 0.03 | [0.00; 0.14] | 0.7% |
| Rompin (Zarith et al., 2022) | 1 | 35 | 0.03 | [0.00; 0.15] | 0.7% |
| Kuala Selangor (Zarith et al., 2022) | 1 | 28 | 0.04 | [0.00; 0.18] | 0.7% |
| Port Dickson (Zarith et al., 2022) | 1 | 26 | 0.04 | [0.00; 0.20] | 0.7% |
| Kuala Kangsar, Perak (Amir et al., 2020) | 1 | 26 | 0.04 | [0.00; 0.20] | 0.7% |
| Kalantan (Saleh Huddin et al., 2019) | 1 | 22 | 0.05 | [0.00; 0.23] | 0.7% |
| Kelantan (Yusuf et al., 2022) | 1 | 22 | 0.05 | [0.00; 0.23] | 0.7% |
| Terengganu (Saleh Huddin et al., 2019) | 4 | 58 | 0.07 | [0.02; 0.17] | 0.9% |
| Batu Pahat (Zarith et al., 2022) | 7 | 93 | 0.08 | [0.03; 0.15] | 0.9% |
| Kangar (Zarith et al., 2022) | 3 | 37 | 0.08 | [0.02; 0.22] | 0.8% |
| Terengganu (Yusuf et al., 2022) | 5 | 58 | 0.09 | [0.03; 0.19] | 0.9% |
| Kota Tinggi (Zarith et al., 2022) | 4 | 46 | 0.09 | [0.02; 0.21] | 0.9% |
| Setiu (Zarith et al., 2022) | 1 | 11 | 0.09 | [0.00; 0.41] | 0.6% |
| Pendang (Zarith et al., 2022) | 2 | 22 | 0.09 | [0.01; 0.29] | 0.8% |
| Jasin (Zarith et al., 2022) | 3 | 32 | 0.09 | [0.02; 0.25] | 0.8% |
| Petaling (Zarith et al., 2022) | 23 | 245 | 0.09 | [0.06; 0.14] | 1.0% |
| Kluang (Zarith et al., 2022) | 1 | 10 | 0.10 | [0.00; 0.45] | 0.6% |
| Pontian (Zarith et al., 2022) | 1 | 9 | 0.11 | [0.00; 0.48] | 0.6% |
| Hulu Langat (Zarith et al., 2022) | 3 | 25 | 0.12 | [0.03; 0.31] | 0.8% |
| Kuala Lipis (Vythilingam et al., 2008) | 10 | 75 | 0.13 | [0.07; 0.23] | 0.9% |
| Temerloh (Zarith et al., 2022) | 9 | 62 | 0.15 | [0.07; 0.26] | 0.9% |
| Seremban (Zamzuri et al., 2020) | 6 | 41 | 0.15 | [0.06; 0.29] | 0.9% |
| Gombak (Zarith et al., 2022) | 2 | 13 | 0.15 | [0.02; 0.45] | 0.8% |
| Kuala Muda (Zarith et al., 2022) | 12 | 66 | 0.18 | [0.10; 0.30] | 0.9% |
| Marang (Zarith et al., 2022) | 5 | 26 | 0.19 | [0.07; 0.39] | 0.9% |
| Kuala Lipis (Zarith et al., 2022) | 5 | 25 | 0.20 | [0.07; 0.41] | 0.9% |
| Kinta (Zarith et al., 2022) | 1 | 5 | 0.20 | [0.01; 0.72] | 0.6% |
| Manjung (Zarith et al., 2022) | 1 | 5 | 0.20 | [0.01; 0.72] | 0.6% |
| Kemaman (Zarith et al., 2022) | 4 | 19 | 0.21 | [0.06; 0.46] | 0.9% |
| Kampar (Zarith et al., 2022) | 2 | 9 | 0.22 | [0.03; 0.60] | 0.7% |
| Pahang (Saleh Huddin et al., 2019) | 43 | 188 | 0.23 | [0.17; 0.30] | 1.0% |
| Selangor (Ho et al., 2010) | 25 | 107 | 0.23 | [0.16; 0.33] | 1.0% |
| Gua Musang (Zarith et al., 2022) | 10 | 38 | 0.26 | [0.13; 0.43] | 0.9% |
| Temerloh, Pahang (Amir et al., 2020) | 9 | 34 | 0.26 | [0.13; 0.44] | 0.9% |
| Kuala Pilah (Zarith et al., 2022) | 7 | 26 | 0.27 | [0.12; 0.48] | 0.9% |
| Tanjung Bungah (Zarith et al., 2022) | 2 | 7 | 0.29 | [0.04; 0.71] | 0.7% |
| Selangor (Atker et al., 2015) | 21 | 70 | 0.30 | [0.20; 0.42] | 1.0% |
| Jempol (Zamzuri et al., 2020) | 6 | 20 | 0.30 | [0.12; 0.54] | 0.9% |
| Maran (Zarith et al., 2022) | 6 | 19 | 0.32 | [0.13; 0.57] | 0.9% |
| Pahang (Yusuf et al., 2022) | 60 | 188 | 0.32 | [0.25; 0.39] | 1.0% |
| Tanah Merah (Zarith et al., 2022) | 11 | 33 | 0.33 | [0.18; 0.52] | 0.9% |
| Pulau Jerejak (Zarith et al., 2022) | 4 | 11 | 0.36 | [0.11; 0.69] | 0.8% |
| Jitra (Zarith et al., 2022) | 12 | 31 | 0.39 | [0.22; 0.58] | 0.9% |
| Taiping (Zarith et al., 2022) | 2 | 5 | 0.40 | [0.05; 0.85] | 0.7% |
| Tampin (Zamzuri et al., 2020) | 32 | 80 | 0.40 | [0.29; 0.52] | 1.0% |
| Bera (Zarith et al., 2022) | 7 | 17 | 0.41 | [0.18; 0.67] | 0.9% |

*Figure S6 (cont.)*

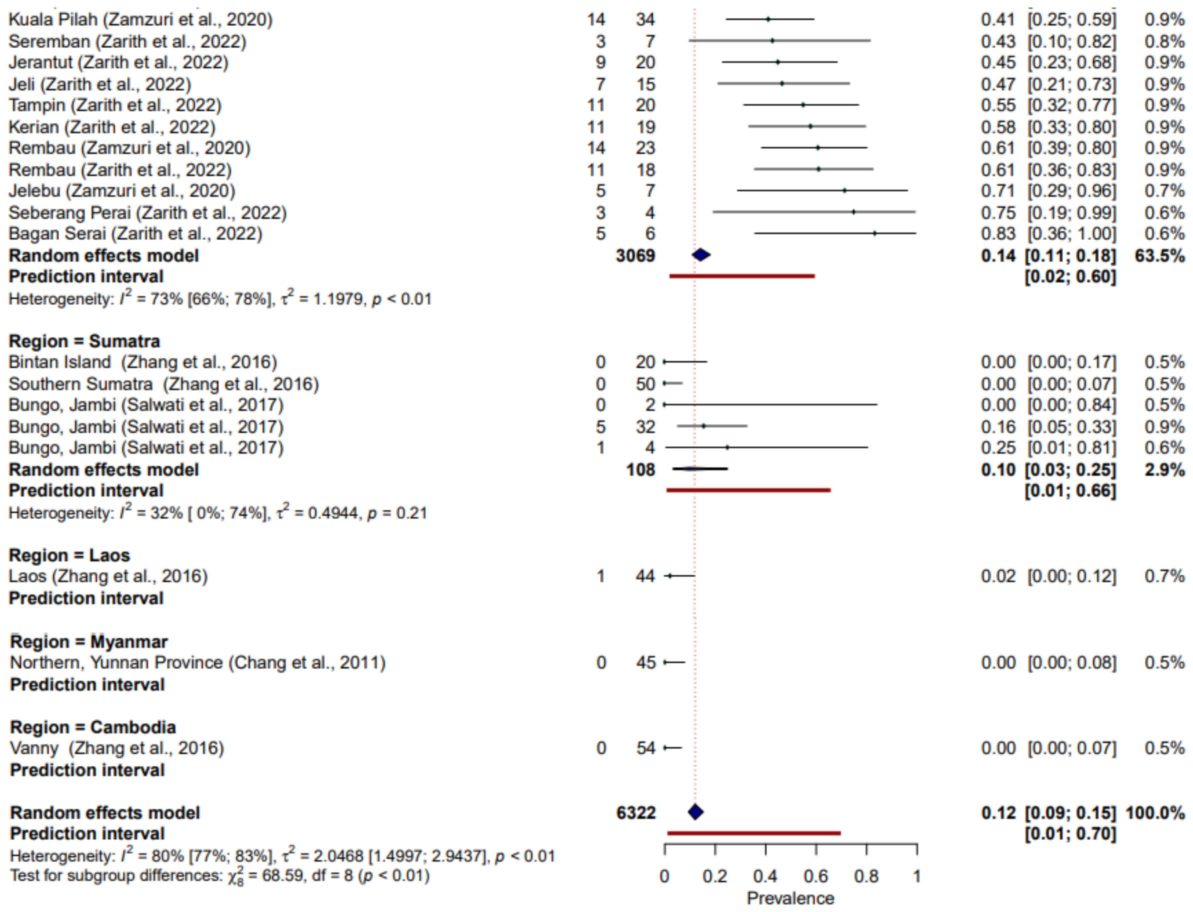

**Figure S6**. *Forest plot of* P. knowlesi *prevalence (%) in all species of NHPs in Southeast Asia, disaggregated by species and sampling site. Random-effects analysis, sub-grouped by region. N=148.*

**APPENDIX D. Remote sensing data and covariate assembly**

Environmental covariates were extracted from satellite-derived remote-sensing datasets. Elevation can be used as a proxy for vector range, with malaria transmission patterns often correspond to altitudinal ranges of vector species[23]. Estimates of human population density provide a measure of urbanisation, used to examine risks related to human settlement proximity. Metrics of deforestation including canopy cover (cumulative loss) and degree of fragmentation, which are key determinants of macaque habitat selection[24] and of mosquito vector breeding sites and were derived from Hansen's Global Forest Watch[25] Land use classification maps derived from the Intact Forest Landscape (IFL)[26] and Copernicus Global Land Service (100m)[27] were also explored to provide more detailed information on specific landscape composition. However, as 77.0% of NHP records were collected before the earliest land classification in 2015 (N=114/148), datasets were of limited utility and not pursued further.

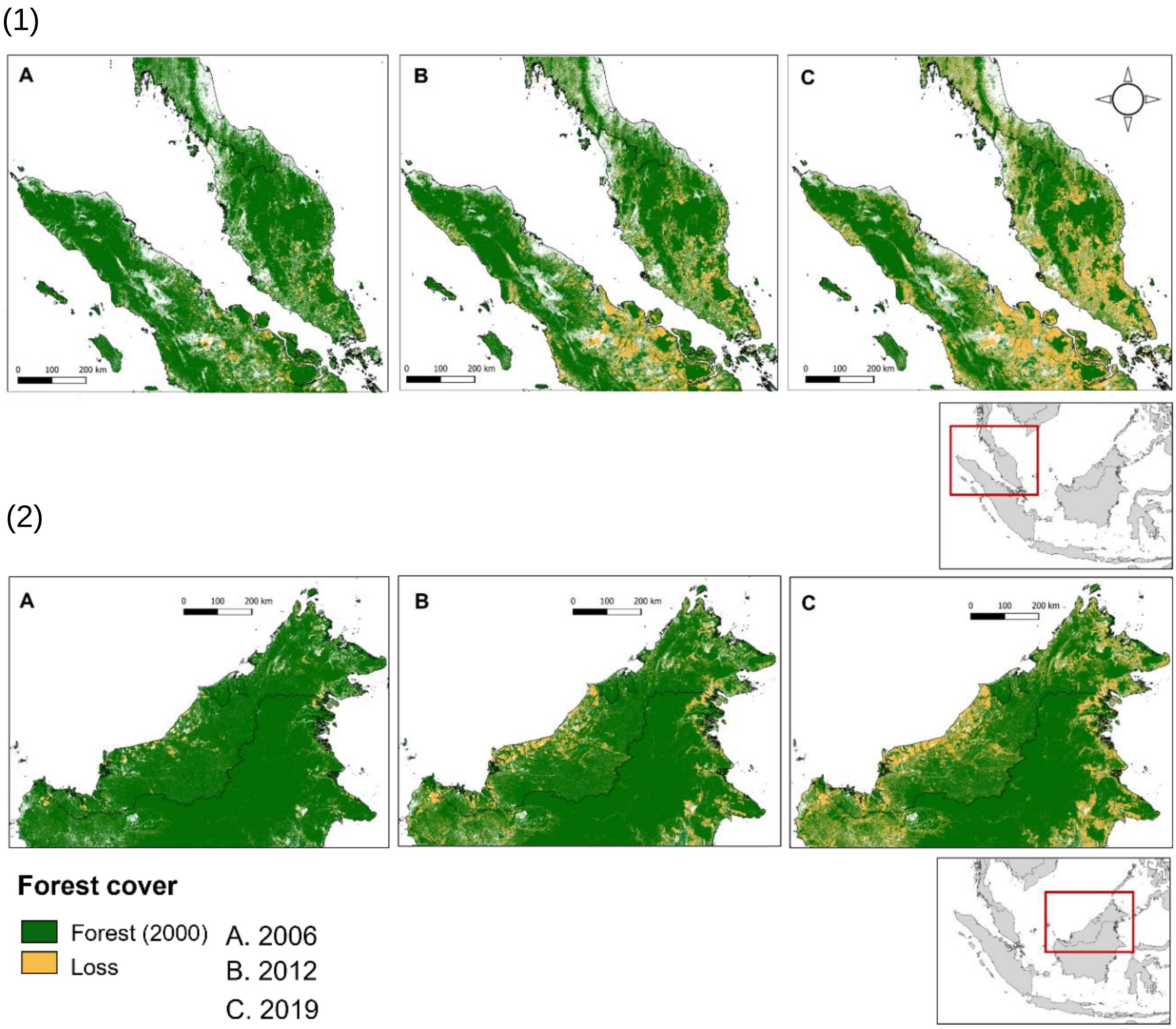

***Figure S7.*** *Recent forest loss in (1) Peninsular Malaysia and (2) Malaysian Borneo, shown for the years 2006/2012/2019*

Gridded UN-adjusted human population estimates were assembled at 1km resolution from WorldPop[28] for multiple timepoints as a measure of urbanisation, a proxy for risks related to human settlement proximity. As minimal variation between timepoints was observed, only 2012 (median year of primate data collection) was retained.

***Table S11.*** *Environmental covariates assembled for regression analysis. Summary values extracted for each covariate within 5, 10 and 20km circular buffers during processing.*

| Covariate | Description | Metric | Resolution | | Processing | Source |
|---|---|---|---|---|---|---|
| | | | Spatial | Temporal | | |
| Population | UN-adjusted gridded posterior population model estimates at 30 arc-seconds resolution | Person count/1km$^2$ | 1km | 2000<br>2012<br>2019 | Population density reclassified as high/low (≤300 persons/km$^2$) in QGIS | WorldPop[28]. Downloaded as tiff files per country in AOI for years 2000/2012/2019 |
| Elevation | Mean height above sea level | m | 1km | 2003 | Mean and SD of continuous elevation per radii. Mean-centred and scaled. Categorised into discrete classifications: low (≤200m), moderate (200–500m) or high elevation (>500m) | NASA SRTM 90m Digital Elevation Database v4.1 (CGIAR-CSI)[29]. Downloaded as a tiff file at 1km resampled resolution |
| Forest | Percentage canopy cover per grid cell. Derived from tree cover (vegetation >5m) and loss (forested to non-forested) | 0–1 | 30m | Annual 2006–2020 | Tree cover classified as ≥ 50% crown density per raster cell, generating binary raster (1=forest, 0=non-forest). Annual cover calculated by subtracting cumulative loss per year 2006–2019. Data records matched to reclassified tile by geolocation and year. Posterior proportions categorised as high (>50%) medium (20–50%) or low (≤20%) | Hansen's Global Forest Watch, 30m resolution Landsat imagery[25]. Tiles downloaded as tiff files for each year 2006–2019 to cover AOI |
| Fragmentation (perimeter: area ratio, PARA) | Perimeter length (m) to patch area (m$^2$) ratio for contiguous forest cover[24] within buffer | PARA>0 | 30m | Annual 2006–2020 | Extracted from annual reclassified tree cover rasters within 5, 10 and 20km circular buffers Output categorised into quartiles | Hansen's Global Forest Watch[25] (as above) |

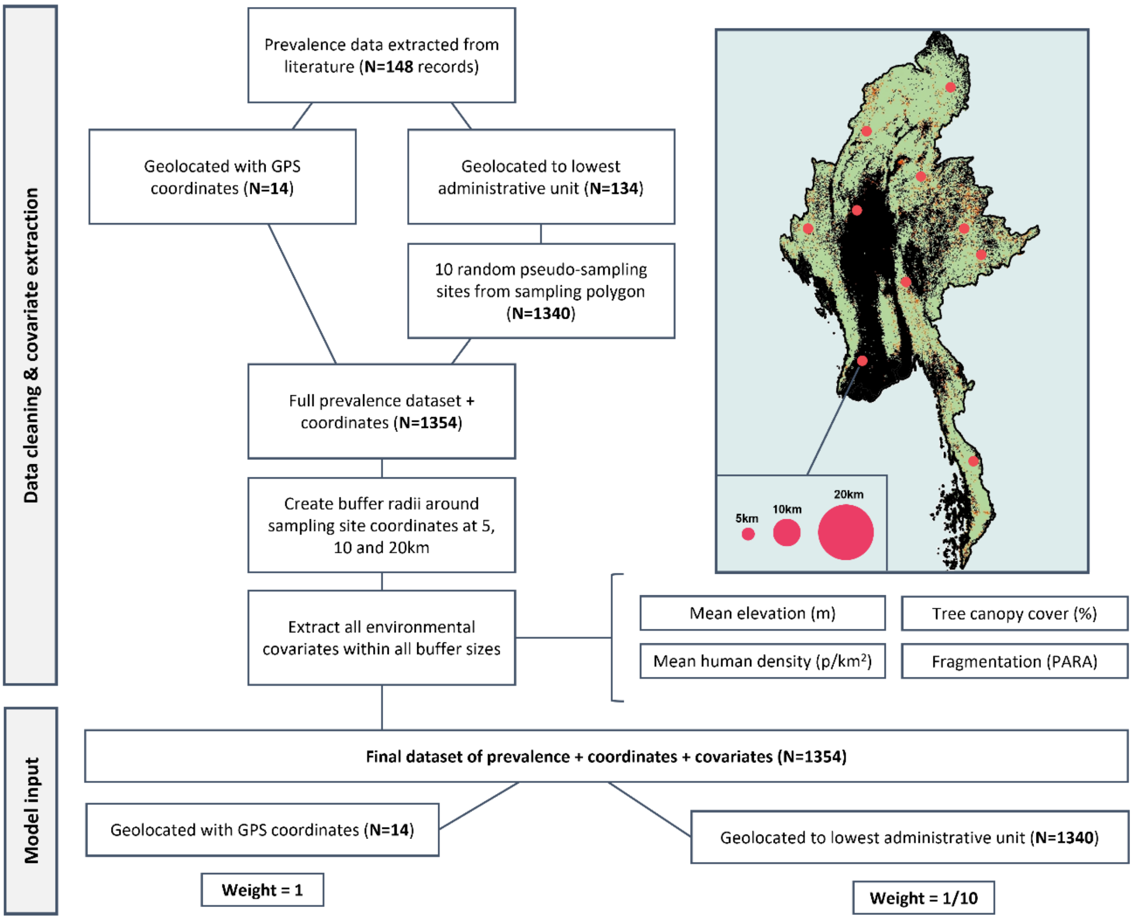

***Figure S8.*** *Flowchart of data processing. Details of pseudo-sampling and environmental covariate extraction at multiple spatial scales to create final weighted dataset (N=1354).*

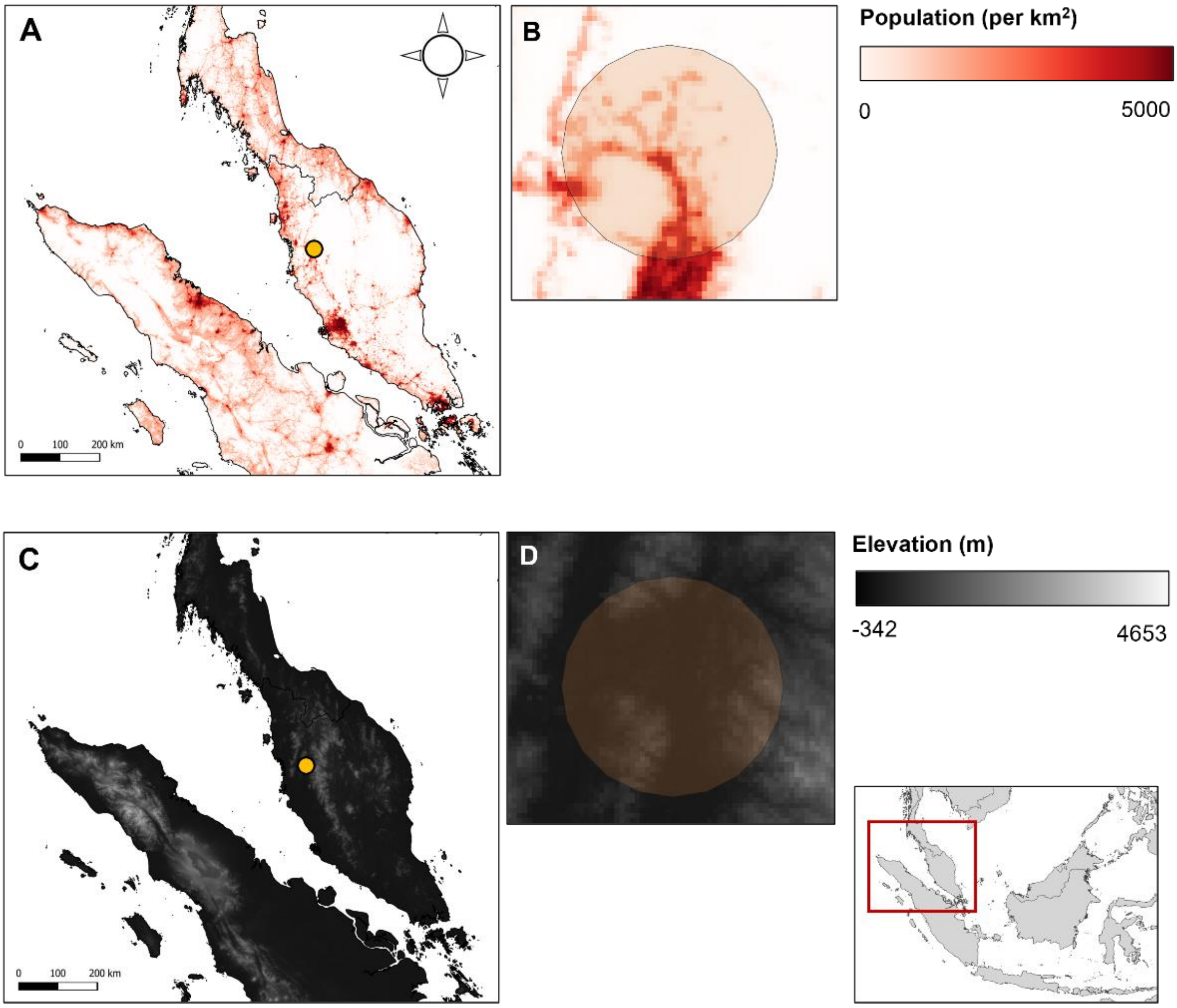

***Figure S9.*** *Example covariate resolutions in Peninsular Malaysia A) point over population density, 1km resolution B) 20km buffer C) data point over SRTM elevation, 1km resolution D) 20km buffer*

Elevation, population density and forest cover all varied markedly across surveyed sites. Forest cover ranges from negligible to near total cover within 5km, and up to 99.96% and 99.64% within 10km and 20km respectively (Table S11, Figure S10). Within a 20km buffer, 46.1% of sites have dense forest cover ≥50% (n=683/1480) and 83.85% have moderate or high forest cover (≥20%), with similar distributions over 5km and 10km. Example buffers over forest cover data can be visualised in Figure S11.

***Table S12.*** *Summary of forest cover data (N=1480)*

| | Mean | SD | Range |
|---|---|---|---|
| Forest cover (5km) | 50.20% | ± 29.29% | 0.00 – 100.00% |
| Forest cover (10km) | 49.68% | ± 27.30% | 0.00 – 99.96% |
| Forest cover (20km) | 48.29% | ± 25.35% | 0.00 – 99.64% |
| Total | | | 1480 (100%) |

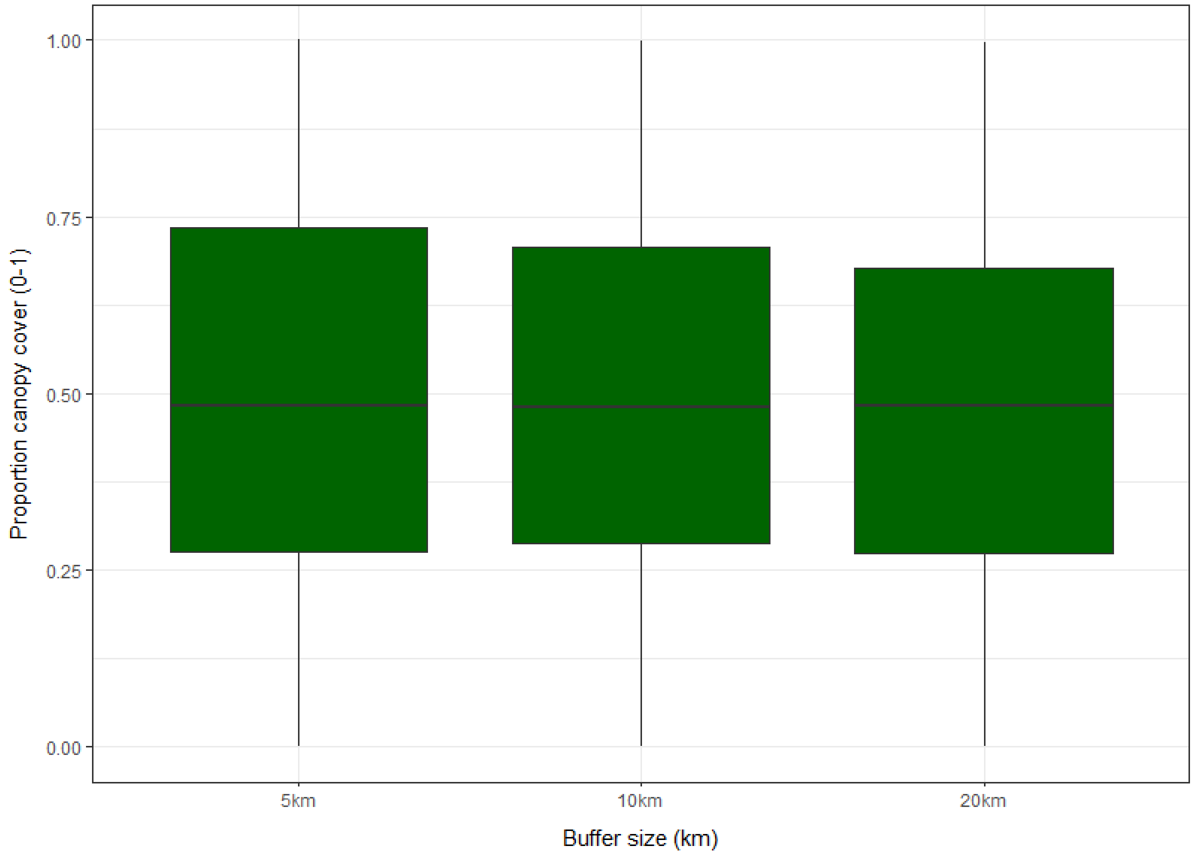

***Figure S10.*** *Boxplots showing distribution and interquartile range (IQR) of proportional forest cover (0-1) for sampling sites within 5, 10 and 20km circular buffers across all sites (N=1354).*

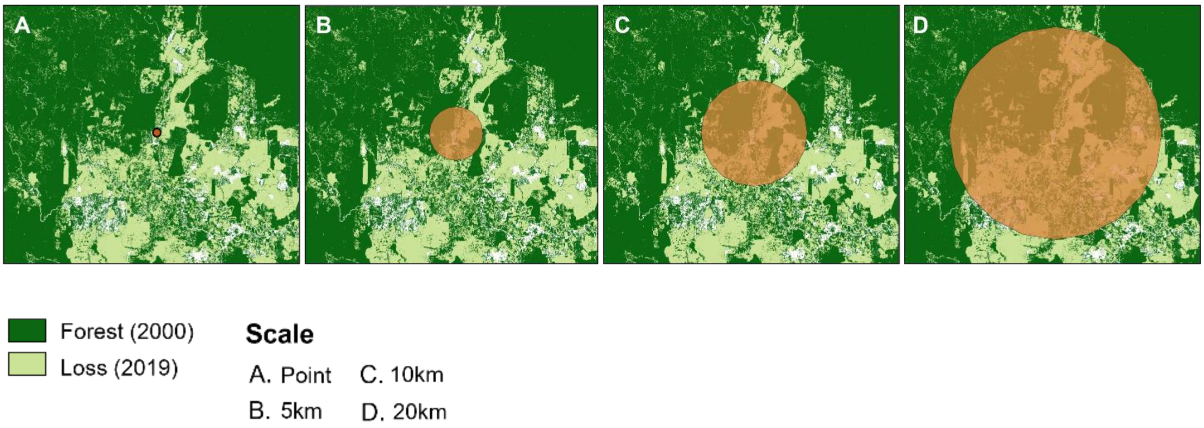

***Figure S11.*** *Examples of buffer zones around macaque sample sites. Shown over forest cover for 2019*[25]

### APPENDIX *E*. Spatial uncertainty

Available spatial resolution of the survey sites varied. 14 records (9.5%, n=14/148) could be geolocated to a point, using geographic coordinates provided or inferred. The remaining 134 were geolocated to the lowest administrative polygon according to GADM boundary definitions (Table S7).

***Table S7.*** *Geo-positioning of available primate survey data*

| | Resolution/GADM* | Records/n | Primates/N | Min. (km²)† | Max. (km²) |
|---|---|---|---|---|---|
| Polygon | Country/GID0 | 6 (4.9%) | 853 (17.3%) | 700 | 77,650 |
| | State/GID1 | 40 (22.0%) | 2699 (32.2%) | 130 | 87,860 |
| | District/GID2 | 88 (61.8%) | 2433 (43.6%) | 270 | 15,890 |
| Point | GCS‡ | 14 (11.4%) | 337 (6.8%) | – | – |
| Total | | 148 (100%) | 4931 (100%) | | |

***Administrative boundaries, as classified by GADM (v3.6)**
†Minimum and maximum size (km²) of polygons containing *P. knowlesi* data at each admin level
‡ Geographic Coordinate System

Crude sensitivity analyses were initially conducted to evaluate use of centroids vs random points to approximate macaque survey site. Using GADM classifications, the largest polygon containing NHP data was identified at each administrative level. 10 points were randomly generated within each polygon in QGIS, with buffers at 5/10/20km. Proportion of forest cover per buffer was extracted, categorised and compared to the forest cover for the centroid (Table S9).

***Table S8.*** *Sensitivity analysis comparing centroid forest cover to 10 randomly generated points, shown per radius for the largest polygon at each GADM level*

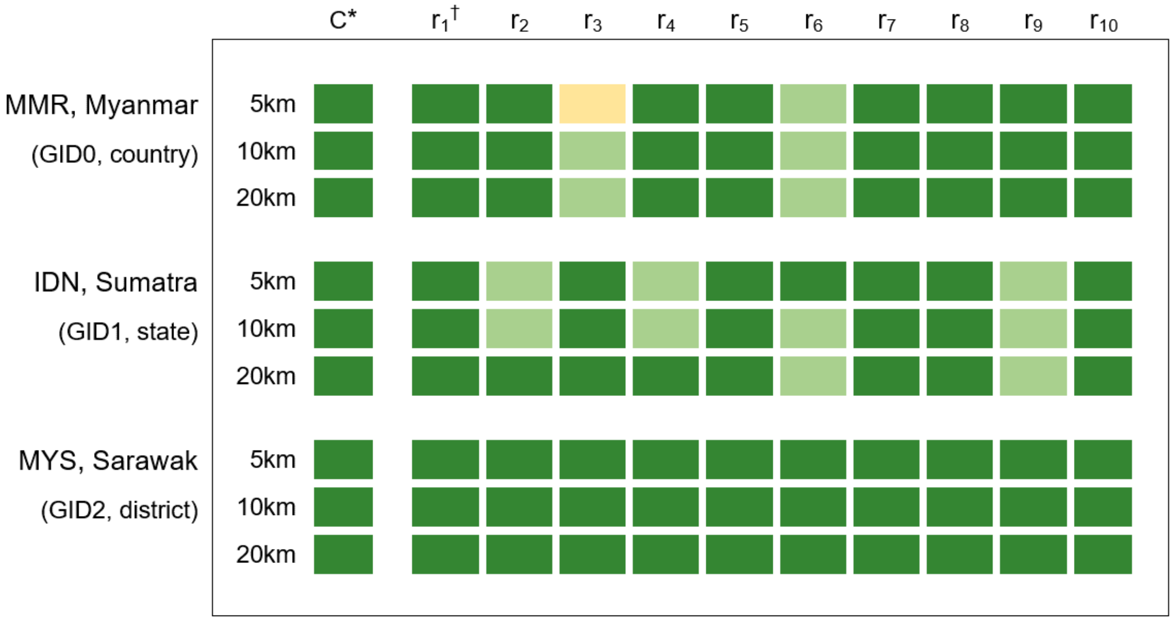

*Centroid of polygon
†Random points ($r_{1-10}$) within polygon
‡Proportion of buffer covered by forest (0<P<1)

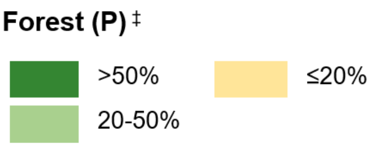

Results indicate that at district level (GID2), minimal change in forest cover between the points was observable (5km: 0.68–0.96). However, for both the state of Southern Sumatra (GID1 5km: 0.26–0.97) and for Myanmar (GID0 5km: 0.16–0.99) variation was observed, with several points classifying as ‘moderate’ (20–50% cover) rather than ‘high’ (>50%) as suggested by the centroid. Overall, results show a disparity between covariates obtained at a central point compared with random points within larger administrative polygons, indicating inadequate sensitivity of this centroids as a proxy for local landscape variables where there is spatial uncertainty.[30]

**A.**

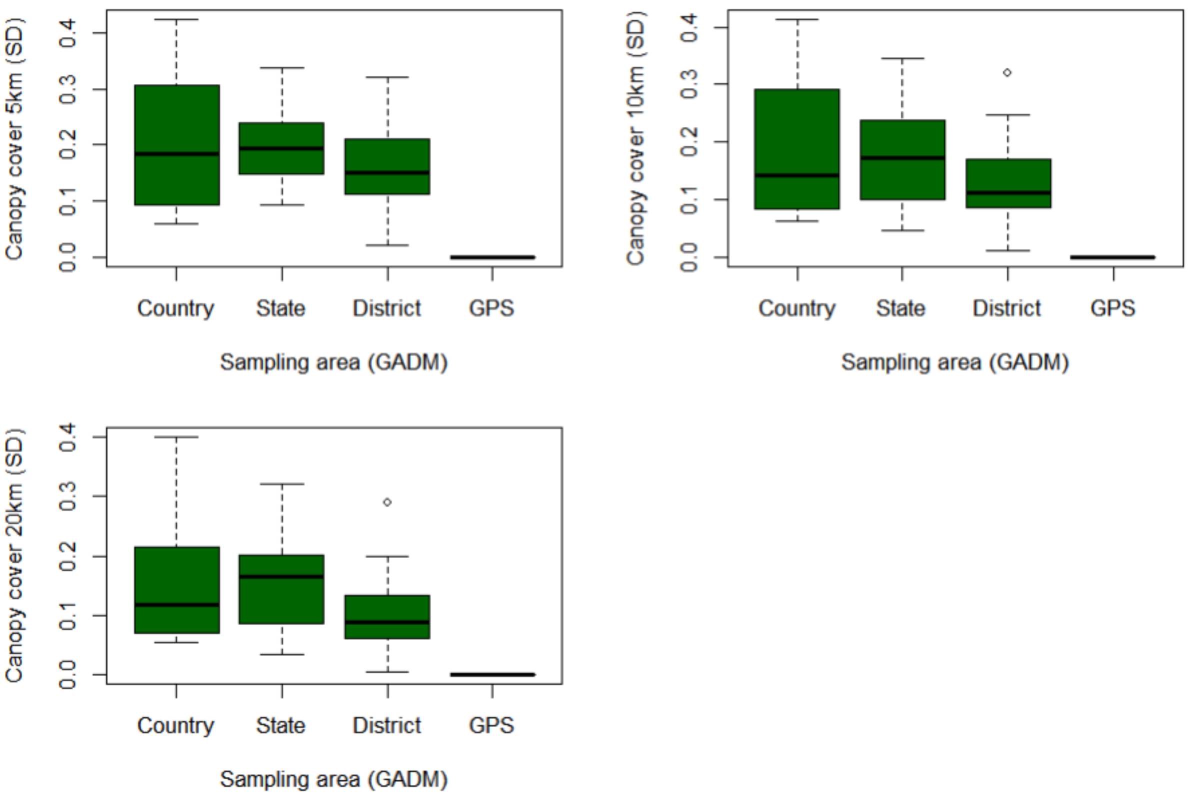
Canopy cover 5km (SD)
Canopy cover 10km (SD)
Canopy cover 20km (SD)
0.0
0.1
0.2
0.3
0.4
Country
State
District
GPS
Sampling area (GADM)

**B.**

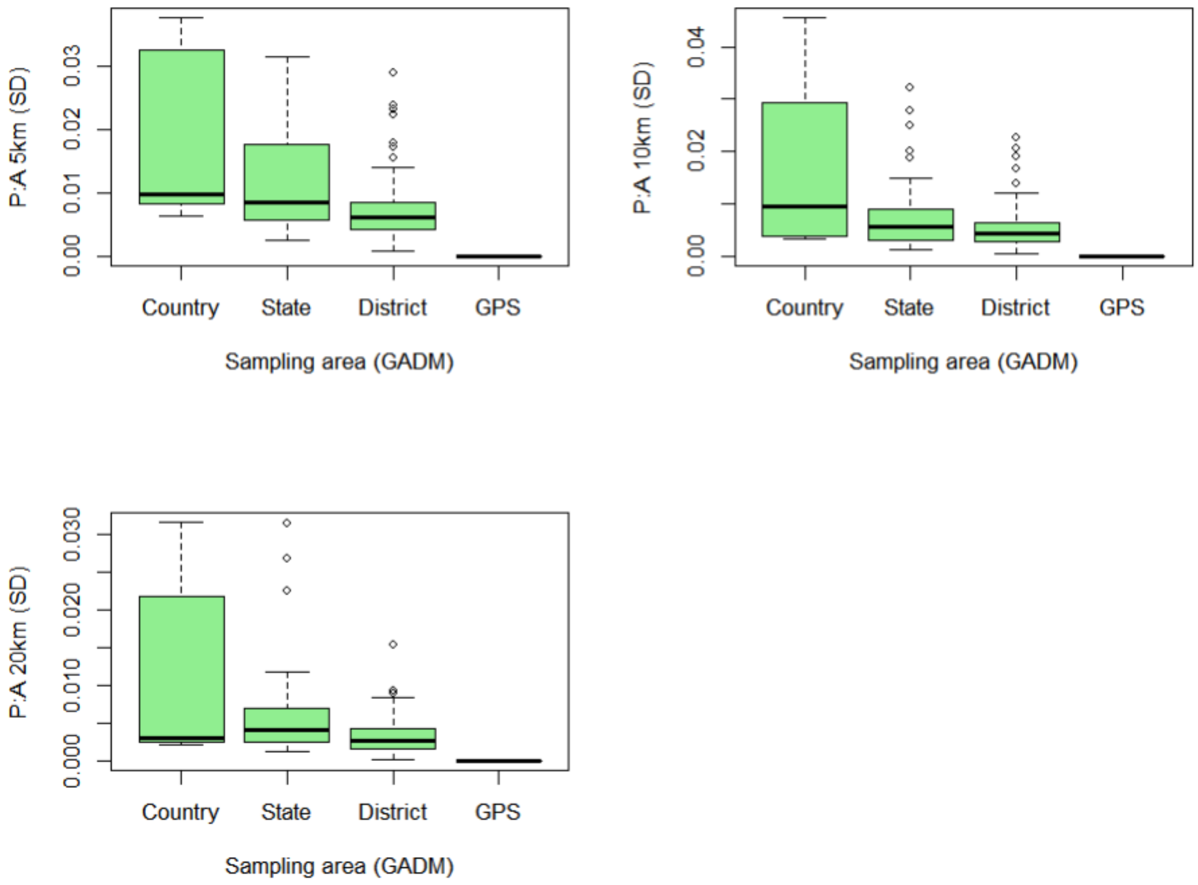
P:A 5km (SD)
P:A 10km (SD)
P:A 20km (SD)
0.00
0.01
0.02
0.03
0.04
0.000
0.010
0.020
0.030
Country
State
District
GPS
Sampling area (GADM)

**C.**

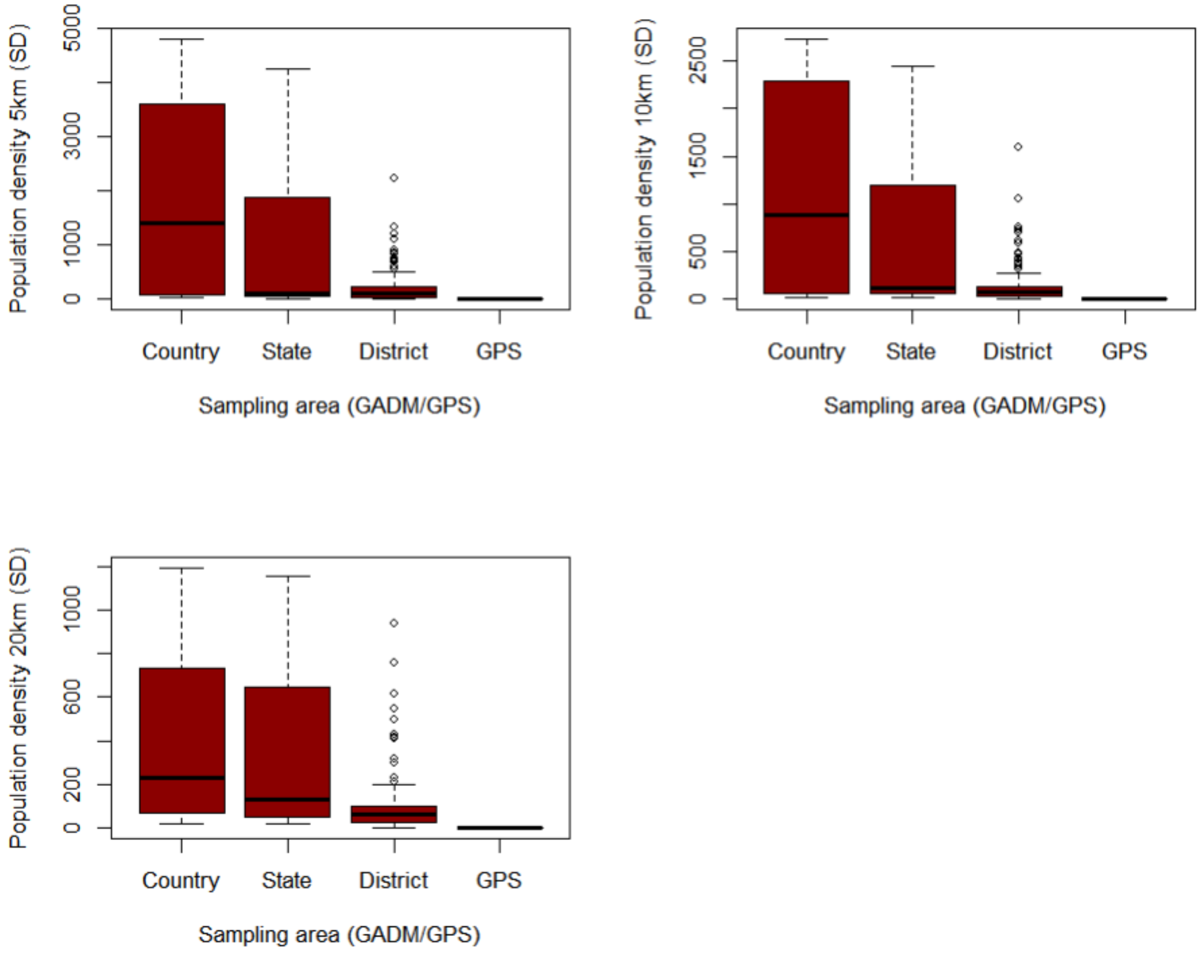

***Figure S12.*** *Standard deviation of environmental covariates across 10 sampling site realisations within 5/10/20km buffers, grouped by administrative boundary size or GPS coordinate* ***A.*** *canopy cover (%)* ***B.*** *forest fragmentation (P: A ratio)* ***C.*** *human population density (p/km$^2$)*

### APPENDIX F. Regression analysis

**Table S13.** Bivariable analysis for *P. knowlesi* in NHP against all covariates at all spatial scales (N=1354).

| | Bivariable analysis | | | |
|---|---|---|---|---|
| Variable | Crude OR | CI95% | | *p* value† |
| Elevation (m) * | | | | |
| ≤5km | 1.18 | (1.07–1.28) | | 0.000562 |
| ≤10km | 1.20 | (1.09–1.31) | | 0.0001246 |
| ≤20km | 1.22 | 1.11–1.33) | | 7.23E-05 |
| Human density (p/km$^2$) * | | | | |
| ≤5km | 0.84 | (0.77– 0.92) | | 4.72E-05 |
| ≤10km | 0.75 | (0.68–0.82) | | 2.70E-12 |
| ≤20km | 0.71 | (0.63–0.79) | | 1.37E-12 |
| Forest cover (%) * | | | | |
| ≤5km | 1.34 | (1.21–1.49) | | 1.86E-08 |
| ≤10km | 1.41 | (1.26–1.57) | | 6.51E-10 |
| ≤20km | 1.47 | (1.30–1.67) | | 8.66E-10 |
| Fragmentation (PARA) * | | | | |
| ≤5km | 0.85 | (0.76–0.95) | | 0.003944 |
| ≤10km | 0.69 | (0.60–0.80) | | 1.80E-07 |
| ≤20km | 0.67 | (0.57–0.79) | | 5.14E-07 |
| PARA$^2$ * | | | Quadratic term | |
| ≤5km | 0.69 | (0.60–0.80) | 0.10 | 2.06E-06 |
| ≤10km | 0.64 | (0.55–0.74) | 0.08 | 2.65E-08 |
| ≤20km | 0.67 | (0.57–0.78) | 0.03 | 2.78E-06 |
| Host species | | | | |
| Other | Ref | | | |
| *M. fascicularis* | 2.37 | (1.25–4.60) | | 0.007971 |

* Continuous variable, mean-centred and scaled

†p value derived from Likelihood ratio test (LRT)

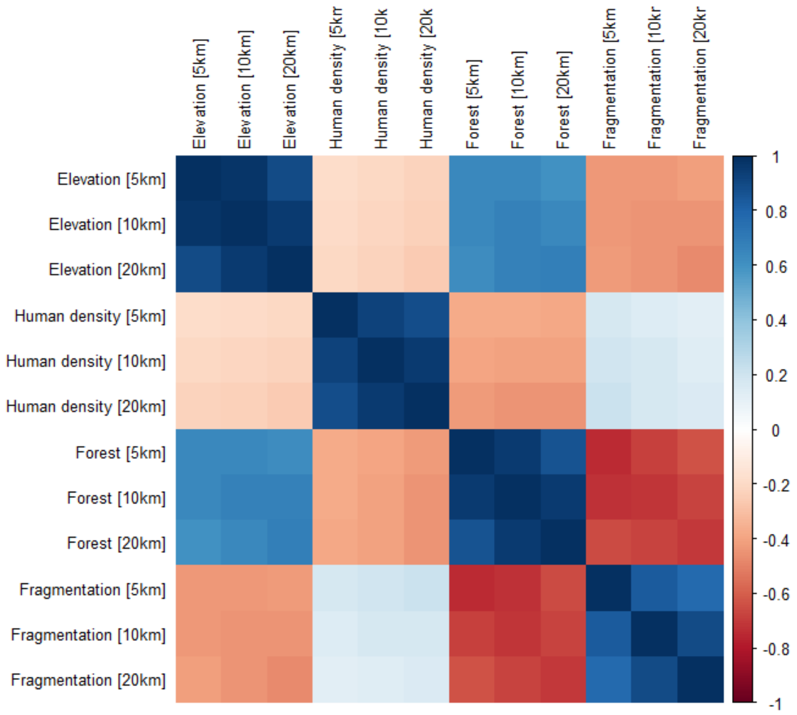

***Figure S13.*** *Spearman's correlation matrix for all candidate covariates at all spatial scales (n=1354).*

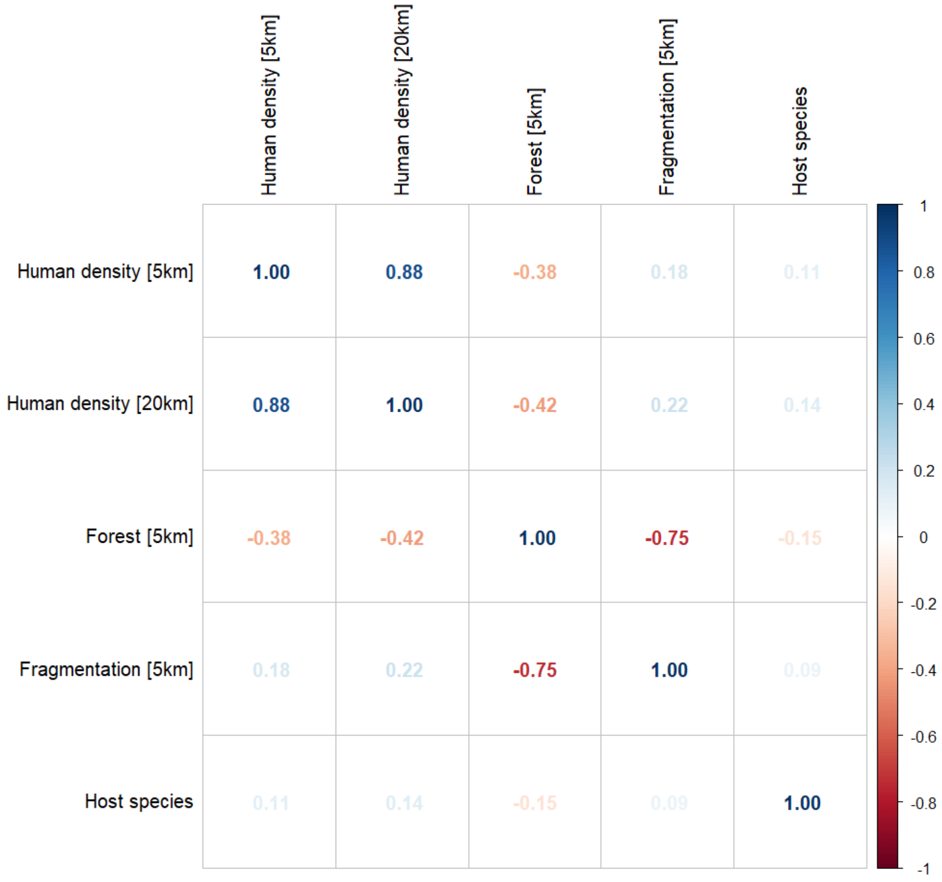

***Figure S14.*** *Spearman's correlation matrix for covariates at selected spatial scales for final model inclusion (n=1354). Percentage forest cover (5km) and forest fragmentation (PARA, 5km) show strong negative correlation (ρ= -0.75).*

**Table S14.** Multivariable binomial regression analysis of *P. knowlesi* prevalence in NHP with environmental covariates at influential spatial scales, full dataset (N=1354). AIC=1229.8.

| Variable | Radius | Multivariable analysis aOR * | CI95% | *p* value † |
|---|---|---|---|---|
| Human density (p/$km^2$) ‡ | | | | |
| | ≤5km | 1.36 | (1.16–1.58) | 1.082E-04 |
| | ≤20km | 0.56 | (0.46–0.67) | 1.311E-10 |
| Forest cover (%)‡ | | | | |
| | ≤5km | 1.38 | (1.19–1.60) | 2.046E-05 |
| Fragmentation (PARA) ‡ | | | | |
| | ≤5km | 1.17 | (1.02–1.34) | 0.0281 |
| Host species | | | | |
| | Other | | Ref | |
| | *M. fascicularis* | 2.50 | (1.31–4.85) | 0.005121 |

*Odds Ratios adjusted for all other variables in the table (aOR). Radius calculated as distance from sample point.

†p value derived from Likelihood ratio test (LRT)

‡Continuous variable, mean-centred and scaled. OR shown per 1 SD increase.

For observations with high geographic uncertainty, random pseudo-sampling of 10 sites (as described in the manuscript and Figure S8) was used to avoid overgeneralisations and biases typical when centroids are used as proxy sampling sites[30], with data weighted accordingly. However, this has the potential to introduce extreme or unrealistic values by generating points in landscapes that are outside reasonable estimations of primate study site. Given this, further sensitivity analyses were conducted to validate the results against geographic precision of observations. Data were first truncated to include only data geolocated to administrative boundaries for relatively small area size (see Figure S12) and exclude highly variable data from country-level boundaries (GID0). Singapore was retained as small administrative unit. GLMM regression models were fit to the truncated dataset.

**Table S15a**. Admin boundary sensitivity analysis. Binomial regression analysis of *P. knowlesi* prevalence in NHP for datapoints assigned to GPS or small sized administrative boundaries (excluding country data) (N=1324).

| **AIC = 1221.9** | | Multivariable analysis | | |
|---|---|---|---|---|
| | | aOR | CI 95% | P value (Wald test) † |
| Human density | [5km] | 1.36 | (1.16–1.58) | *** |
| Human density | [20km] | 0.56 | (0.46–0.67) | *** |
| Forest cover (%) | [5km] | 1.38 | (1.19–1.60) | *** |
| Fragmentation (PARA) | [5km] | 1.18 | (1.02–1.34) | * |
| Host group | Other | REF | | |
| | *M. fascicularis* | 2.51 | (1.–.31–4.85) | ** |

†Signif. codes:  0 '***' 0.001 '**' 0.01 '*' 0.05 '.' 0.1 ' ' 1

Administrative boundaries are arbitrary categories and vary considerably in size and landscape consistency. To better evaluate environmental uncertainty associated with each observation, standard deviation (SD) of the covariate values within each set of 10 environmental realisations was calculated (resulting in a single standard deviation value for each covariate at each scale for every prevalence data point). Studies for which the uncertainty (SD) of covariates exceeded half of the maximum standard deviation were censored to avoid spurious associations derived from unreliable/extreme values for both forest cover (5km) and fragmentation (5km). Regression models were fit to the winsorized dataset and compared to results from the full dataset to ensure that associations are robust.

**Table S16a**. Distribution of standard deviations across 10 environmental covariates per prevalence data point for landscape variables at all spatial scales (N=1354)

| **Covariate** | **Mean** | **Range** | **Median** | **IQR** |
|---|---|---|---|---|
| Canopy [5km] * | 0.1588 | 0.0000-0.4237 | 0.1534 | 0.1039-0.2277 |
| Canopy [10km] | 0.1319 | 0.0000-0.4124 | 0.1177 | 0.0827-0.1891 |
| Canopy [20km] | 0.1051 | 0.0000-0.3999 | 0.0921 | 0.0541-0.1635 |
| Fragmentation [5km] ***** | 0.0083 | 0.0000-0.0375 | 0.0063 | 0.0041-0.0094 |
| Fragmentation [10km] | 0.0061 | 0.0000-0.0455 | 0.0043 | 0.0025-0.0071 |
| Fragmentation [20km] | 0.0041 | 0.0000-0.0316 | 0.0027 | 0.0017-0.0047 |

*Spatial scales selected in final variables

**Table S16b**. Tree canopy cover sensitivity analysis. Binomial regression of *P. knowlesi* prevalence in NHP for datapoints, with data where SD > ½ the maximum for tree canopy within 5km (N=814).

| **AIC = 771.1** | | Multivariable analysis | | |
|---|---|---|---|---|
| | | aOR | CI 95% | P value (Wald test) † |
| Human density | [5km] | 0.90 | (0.67–1.20) | - |
| Human density | [20km] | 0.72 | (0.50–1.01) | . |
| Forest cover (%) | [5km] | 1.70 | (1.30–2.24) | *** |
| Fragmentation (PARA) | [5km] | 1.38 | (1.01–1.88) | * |
| Host group | Other | REF | | |
| | *M. fascicularis* | 2.63 | (1.35–5.21) | ** |

†Signif. codes:  0 '***' 0.001 '**' 0.01 '*' 0.05 '.' 0.1 '-' 1

**Table S16c**. Landscape fragmentation sensitivity analysis. Binomial regression of *P. knowlesi* prevalence in NHP for datapoints, with datapoints where SD > ½ the maximum for fragmentation within 5km (N=1134).

| **AIC = 982.9** | | Multivariable analysis | | |
|---|---|---|---|---|
| | | aOR | CI 95% | P value (Wald test) † |
| Human density | [5km] | 0.91 | (0.69–1.18) | - |
| Human density | [20km] | 0.69 | (0.52–0.93) | * |
| Forest cover (%) | [5km] | 1.31 | (1.08–1.60) | ** |
| Fragmentation (PARA) | [5km] | 1.18 | (0.90–1.54) | - |
| Host group | Other | REF | | |
| | *M. fascicularis* | 2.52 | (1.33–4.87) | ** |

†Signif. codes:  0 '***' 0.001 '**' 0.01 '*' 0.05 '.' 0.1 ' ' 1

Sensitivity analysis has shown that the trends are robust when the data is constrained according to small administrative boundaries or by measures of spatial uncertainty in the environmental variables. However, given that a proportion of points are randomly generated, questions remain about how suitable the resulting sites are for macaque species and consequently whether the associations observed are a realistic indication of ecological trends. To address this, points were subset according to macaque species habitat suitability maps, derived from Moyes et al (2016)[31].

Predicted occurrence maps were combined for three species *Macaca fascicularis, Macaca nemestrina* and *Macaca leonina* to create a joint macaque extent for Southeast Asia. Binary maps of predicted habitat extent were then generated using thresholds of moderate (predictions of 0.5 and above), high (>0.75) and very high predicted probability of occurrence (>0.9)[31] (Figure S15a–c). Datapoints from the main analysis were then overlayed with each map, and any points (± 5km buffer) that occurred outside of predicted habitat extent for macaque species were removed. Regression models were then fit to the reduced datasets to assess whether associations observed are plausible according to macaque ecology (Table S17a–c).

**Figure S15a.** Distribution and habitat range of dominant macaque species (*M. fascicularis, M. nemestrina, M. leonina)* according to predicted probability of occurrence ≥ 0.5 (on a scale of 0 to 1.0) per 5x5km pixel.

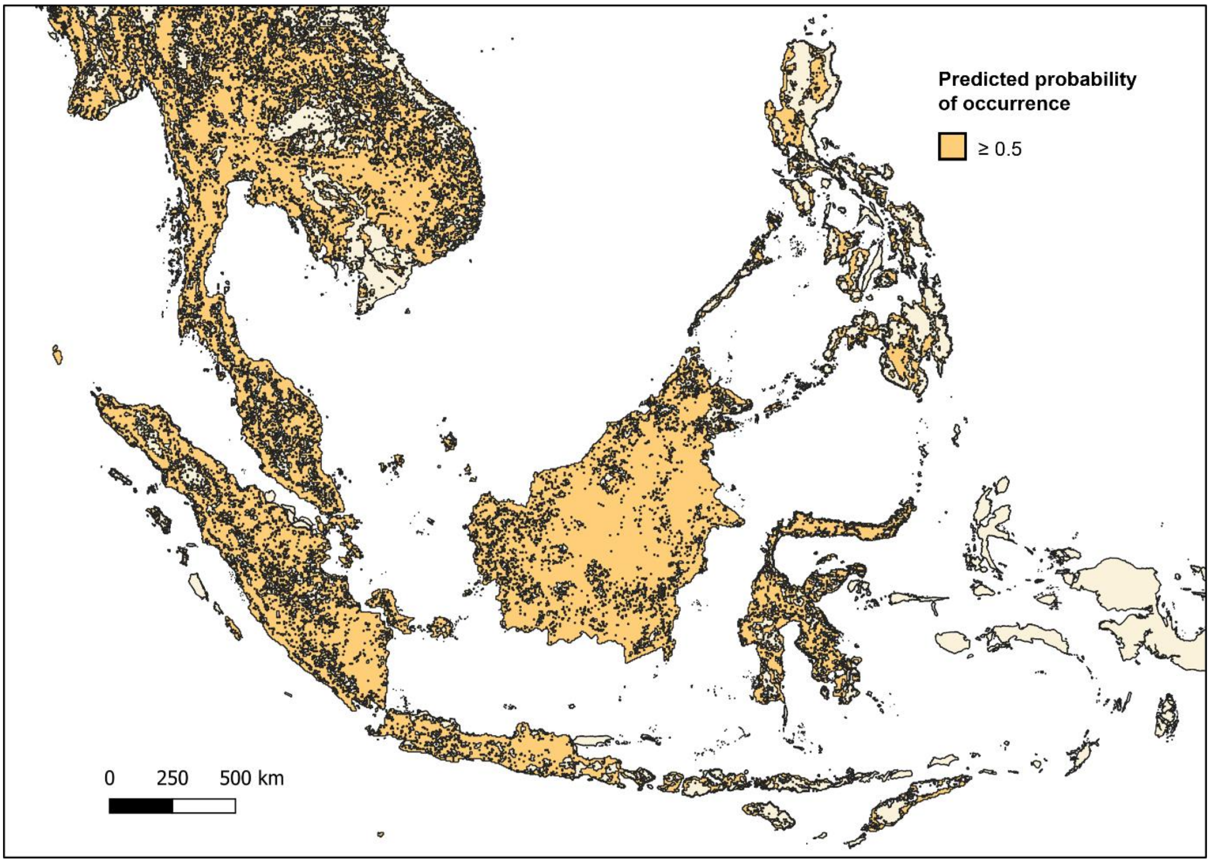

**Table S17a**. Macaque habitat suitability sensitivity analysis. Binomial regression of *P. knowlesi* prevalence in NHP for datapoints, including only datapoints with 5km buffers that intersect with areas with ≥ 0.5 probability of predicted macaque occurrence (N=1331).

| **AIC = 1197.2** | | Multivariable analysis | | |
|---|---|---|---|---|
| | | aOR | CI 95% | P value (Wald test) † |
| Human density | [5km] | 1.32 | (1.13–1.54) | *** |
| Human density | [20km] | 0.55 | (0.45–0.66) | *** |
| Forest cover (%) | [5km] | 1.30 | (1.12–1.52) | *** |
| Fragmentation (PARA) | [5km] | 1.12 | (0.97–1.29) | - |
| Host group | Other | REF | | |
| | *M. fascicularis* | 2.48 | (1.31–4.82) | ** |

†Signif. codes: 0 '***' 0.001 '**' 0.01 '*' 0.05 '.' 0.1 ' ' 1

**Figure S15b.** Distribution and habitat range of dominant macaque species (*M. fascicularis, M. nemestrina, M. leonina)* according to predicted probability of occurrence ≥ 0.75 (on a scale of 0 to 1.0) per 5x5km pixel.

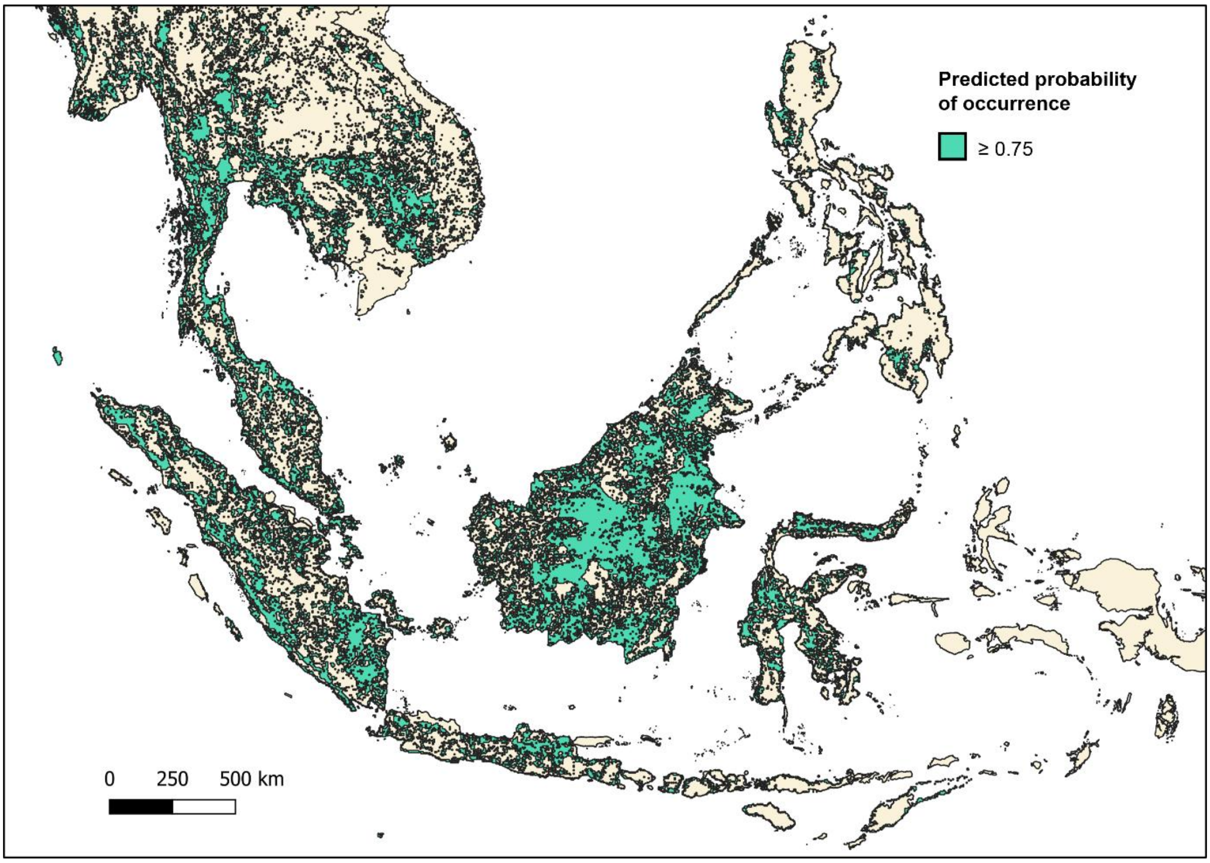

**Table S17b**. Macaque habitat suitability sensitivity analysis. Binomial regression of *P. knowlesi* prevalence in NHP for datapoints, including only datapoints with 5km buffers that intersect with areas with ≥ 0.75 probability of predicted macaque occurrence (N=1177).

| **AIC = 1115.5** | | Multivariable analysis | | |
|---|---|---|---|---|
| | | aOR | CI 95% | P value (Wald test) † |
| Human density | [5km] | 1.34 | (1.14–1.58) | *** |
| Human density | [20km] | 0.57 | (0.47–0.69) | *** |
| Forest cover (%) | [5km] | 1.23 | (1.04–1.47) | * |
| Fragmentation (PARA) | [5km] | 1.04 | (0.86–1.24) | - |
| Host group | Other | REF | | |
| | *M. fascicularis* | 2.69 | (1.38–5.38) | ** |

†Signif. codes: 0 ‘***’ 0.001 ‘**’ 0.01 ‘*’ 0.05 ‘.’ 0.1 ‘ ’ 1

**Figure S15c.** Predicted distribution and habitat range of all macaque species (*M. fascicularis, M. nemestrina, M. leonina)* according to predicted probability of occurrence ≥ 0.9 (on a scale of 0 to 1.0) per 5x5km pixel.

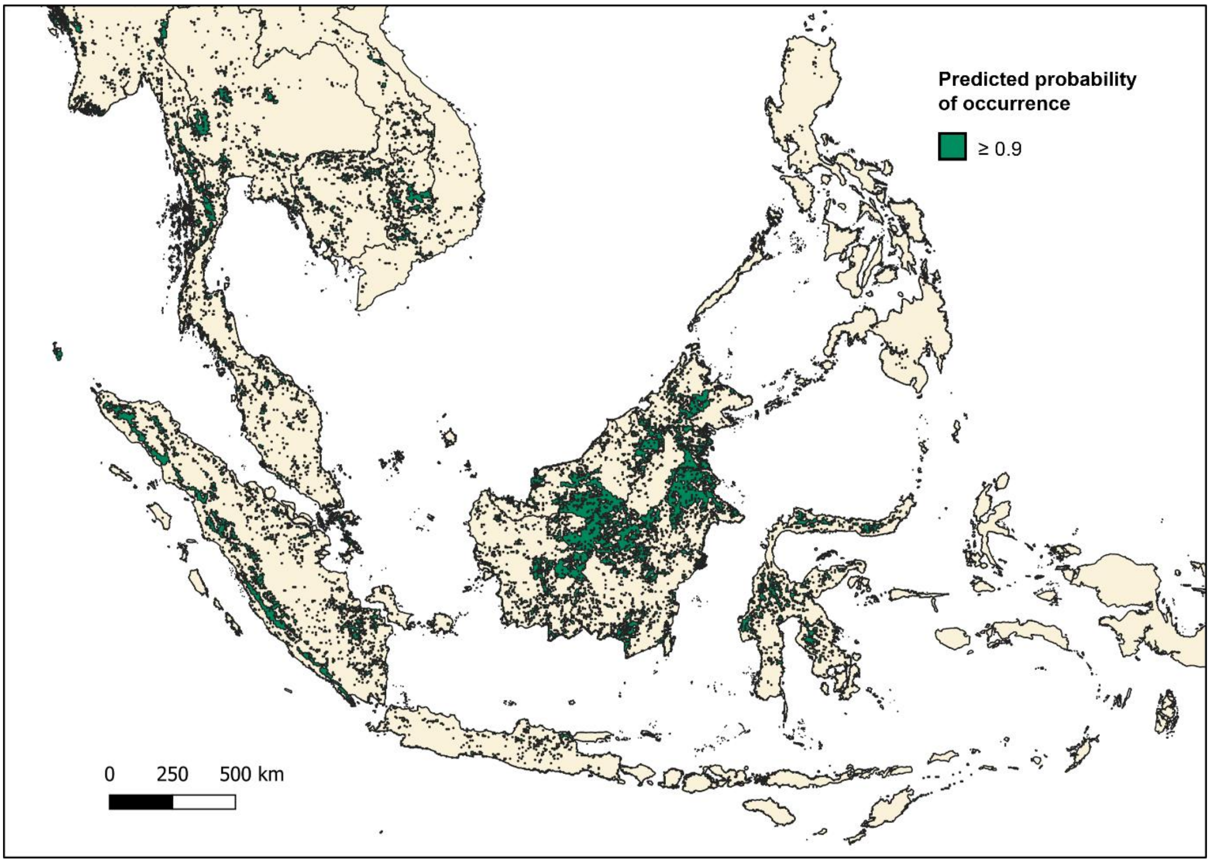

**Table S17c.** Macaque habitat suitability sensitivity analysis. Binomial regression of *P. knowlesi* prevalence in NHP for datapoints, including only datapoints with 5km buffers that intersect with areas with ≥ 0.9 probability of predicted macaque occurrence (N=567).

| **AIC = 685.2** | | Multivariable analysis | | |
|---|---|---|---|---|
| | | aOR | CI 95% | P value (Wald test) † |
| Human density | [5km] | 1.86 | (1.49–2.32) | *** |
| Human density | [20km] | 0.36 | (0.26–0.49) | *** |
| Forest cover (%) | [5km] | 1.47 | (1.14–1.90) | ** |
| Fragmentation (PARA) | [5km] | 1.35 | (1.02–1.77) | * |
| Host group | Other | REF | | |
| | *M. fascicularis* | 3.13 | (1.50–6.75) | ** |

†Signif. codes: 0 ‘***’ 0.001 ‘**’ 0.01 ‘*’ 0.05 ‘.’ 0.1 ‘ ’ 1

15. Unpublished. Personal Correspondance. Published online 2013.

16. Unpublished. Personal Correspondance. Published online 2015.

29. Jarvis, A., H.I. Reuter, A. Nelson EG. Hole-filled SRTM for the globe Version 4, available from the CGIAR-CSI SRTM 90m Database (http://srtm.csi.cgiar.org).

30. Cheng Y, Tjaden NB, Jaeschke A, Thomas SM, Beierkuhnlein C. Using centroids of spatial units in ecological niche modelling: Effects on model performance in the context of environmental data grain size. *Global Ecology and Biogeography*. 2021;30(3):611-621. doi:https://doi.org/10.1111/geb.13240
